# Right anterior insula effective connectivity impairs intrinsic BOLD fluctuations in dorsal attention network in adolescents and young adults with borderline personality symptoms

**DOI:** 10.1101/2022.08.08.503183

**Authors:** Nathan T. Hall, Michael N. Hallquist

**Author notes:** Corresponding author.* Michael Hallquist, Department of Psychology and Neuroscience, University of North Carolina at Chapel Hill. 235 E. Cameron Avenue, Chapel Hill, NC 27599. Disclosures.* The authors have no financial interests to disclose.

## Abstract

**Background:** Borderline Personality Disorder (BPD) symptoms often emerge in adolescence. However, little is known about the functional organization of intrinsic brain networks in young people with BPD symptoms.

**Methods:** In this study we collected resting-state fMRI data in a sample of adolescents and young adults with (n_BPD_ = 40) and without BPD (n_HC_= 42) symptoms. Using a detailed cortico-limbic parcellation coupled with graph theoretical analyses, we tested for group and age-related differences in regional functional and effective connectivity (FC, EC) and amplitude of low frequency fluctuations (ALFF). We conducted a series of analyses that progressed from global network properties to focal tests of EC amongst nodes in Salience (SN) and Dorsal Attention Networks (DAN).

**Results:** At the regional level, regularized regression analyses revealed a broad pattern of hyper-connectivity and heightened ALFF in R dorsal anterior insula (daIns), in addition to hypoconnectivity in R temporal-parietal junction (TPJ) and decreased ALFF in multiple DAN regions. Furthermore, analyses of EC amongst daIns, TPJ, and DAN revealed that in BPD participants daIns exerts a heightened influence on TPJ and DAN regions. Finally, multivariate mediation models indicated that lower DAN_ALFF_ was differentially predicted by EC from TPJ and daIns.

**Conclusions:** Our findings provide converging evidence that heightened EC from daIns impairs network-wide ALFF in DAN both directly and indirectly via impaired TPJ functioning. We interpret this pattern of findings in line with an “attentional hijacking” account of borderline personality.

---

Symptoms of Borderline Personality Disorder (BPD) such as emotion dysregulation and nonsuicidal self-injury typically emerge in adolescence (1,2). Although the maturation of circuits underlying emotion regulation and cognitive control is remarkable during this period (3), little is known about macroscale functional brain organization in young people with BPD symptoms. The current study investigated interactions among intrinsic connectivity networks in adolescents and young adults with BPD symptoms using three major BOLD indices of resting-state brain function: functional and effective connectivity (FC and EC, respectively) and regional amplitude of low-frequency fluctuations (ALFF; 4–6). Through a series of analyses that progress from broad network properties to focal tests of the relation between ALFF and EC in specific networks, we show that heightened resting-state effective connectivity of the R dorsal anterior insula (daIns) in BPD impairs ALFF in the Dorsal Attention Network (DAN).

Our study builds on evidence from developmental neuroscience that the organization of the brain into distinct networks (aka “modules”) is well-established by late childhood, though connectivity patterns amongst these networks undergoes a protracted refinement in adolescence (7–9). By late childhood, intrinsic networks have *segregated* (i.e., increased within-network connections), establishing the functional specializations of each network (9–11). However, during adolescence, functional *integration* among networks facilitates widespread communication via “connector hub” regions that serve as highways of between-network communication, which may underlie adolescence-related improvements in behavioral performance (8).

Crucially, in adolescence, the Salience Network (SN; 12,13) undergoes substantial changes in between-network connectivity compared to other intrinsic networks (8,14). SN is composed primarily of dorsal anterior cingulate (dACC) and aIns and plays a key role in detecting and orienting attention to stimuli that are behaviorally relevant or perceptually salient (12,15). A recent taxonomy of intrinsic networks expanded SN to include right temporal-parietal junction (TPJ) and inferior frontal gyrus (IFG) as well as numerous subcortical regions (16). SN is thought to organize dynamic switches among task-positive networks (DAN, Fronto-Parietal Networks) and the task-negative Default Mode Networks (DMN; 15,17–19). In other words, SN monitors both internal and external events and triggers the appropriate networks to come online to implement either higher-order cognitive functions (DAN, FPN) or self-reflective internally directed thought (DMN). Within SN, the insula is a multimodal hub of cognitive-emotional functioning that receives sensory, cognitive, and homeostatic inputs; its integration of these signals facilitates flexible decision-making (13,20–22). More importantly, numerous studies of insula EC converge on its role as a central “outflow hub”; its widespread projections to cortex suggest a unique ability to control macroscale brain dynamics (18,19).

Given SN’s involvement in a wide array of socio-cognitive and emotional processes and its protracted development in adolescence, it has become a key target in understanding the emergence of psychopathology (23–27). However, SN functioning in adolescents and young adults with BPD symptoms is poorly understood. Task-based fMRI studies of BPD have implicated a diverse set of regions involved in cognitive control, emotion regulation, and social cognition (28–32). However, few studies of intrinsic connectivity in adults with BPD have focused on SN (33–35). These studies have provided promising, though mixed, results regarding SN connectivity in borderline personality^1^. For example, two connectivity studies using ICA reported contradictory results: one found greater SN-DMN connectivity in BPD, while the other reported decreased SN-DMN connectivity (33,34).

This contradiction notwithstanding, there is good reason to believe that aberrant segregation and/or integration of DMN, SN, and task-positive networks may be associated with the expression of BPD symptoms. An influential “triple network model” of psychopathology argues that disrupted interactions among these networks likely contribute to the pathophysiology of numerous psychiatric disorders (23). This model posits that aberrant salience signals computed in aIns project to DMN and fronto-parietal regions, with major downstream effects on the cognitive processes implemented by these networks. For example, hyperactivity/connectivity of the aIns has been proposed to support heightened interoceptive prediction signals in anxiety (36), while hypoactivity of the aIns may underlie weakened salience mapping to social stimuli in autism (27,37).

Although SN likely plays an important role in psychopathology (38), the role of SN’s connectivity with other networks and/or aberrant functioning of specific brain regions within SN remain topics of active inquiry in clinical studies. From the perspective of the triple network model, interpersonal hypersensitivity and volatile emotionality in BPD can be understood in terms of heightened salience mapping, with corresponding activity and connectivity of core SN regions. These salience signals would likely disrupt networks involved in planning and goal-directed behavior (FPN) and self-referential thought (DMN). Importantly, this model does not distinguish between task-positive networks involved in cognitive control and attentional control [FPN, DAN respectively; 38]. In a separate line of research, regions in SN have been shown to dynamically interact with parietal regions in DAN by generating attentional reorienting signals that interrupt transient attentional processing in DAN (40). However, aberrant communication between SN and DAN has received little attention in clinical neuroimaging studies.

Studies of SN in BPD have reported broad abnormalities, yet they have not clarified whether aberrant SN function is localized to specific regions, reflects interactions among SN regions, or may be a network-level phenomenon involving SN’s coordination with other networks. Moreover, essentially no prior studies of BPD have characterized the functioning of intrinsic networks within a graph theoretical framework, the preferred analytic approach in network neuroscience (41,42). Graph theory represents networks in terms of regions (‘nodes’) and connections amongst regions (‘edges’) and offers a broad palette of metrics/analytic approaches that quantify an array of network properties. Importantly, these network properties can be conceptualized in terms of telescoping levels of analysis (43), from global (average connection strength across the entire network) to highly specific (edge characteristics between two nodes). Furthermore, while most connectivity studies in BPD focus on FC (undirected/correlational connectivity), modern EC models support inferences about the directionality of connectivity amongst regions (44–46). In a prior report, we investigated BPD-related differences in EC of amygdala subnuclei and targeted regions in mPFC (47), building on evidence of fronto-limbic dysfunction in BPD (28,48,49). Here, we significantly broadened our scope to a whole-brain connectivity analysis, placing a particular emphasis on intrinsic activity and connectivity of SN and its constituent regions.

While connectivity analyses reveal features of the brain’s functional network architecture, they provide less information about the magnitude of intrinsic activity. Markers of intrinsic activity provide information on the functional integrity of synchronized neural activity in a brain region. For example, FDG-PET studies have found a pattern of glucose hypometabolism in the medial PFC in BPD patients, suggesting abnormalities in the functioning of this region (50). In fMRI data, ALFF is a measure of the amplitude of low-frequency BOLD oscillations [0.01-0.1 Hz; 51], approximating the resting-state activity of a region (52). More specifically, basic studies of ALFF suggest a positive linear relationship between ALFF and glucose metabolism (53) and an inverse linear relationship between ALFF and GABA levels (54). Although ALFF provides an indirect index of regional intrinsic activity^2^, to our knowledge no studies have focused on how connectivity between regions or networks may impair or bolster ALFF.

In a sample of adolescents and young adults with and without BPD symptoms, we examined whole-brain intrinsic connectivity using graph theoretical network measures and tested how connectivity patterns amongst these intrinsic networks are related to regional ALFF. As detailed below, our results suggest that in adolescents and young adults with BPD symptoms, the right dorsal aIns (daIns) is hyperactive (heightened ALFF) and exhibits a strengthened directed influence on DAN, which impairs network-level activity in DAN.

## Materials and Methods

### Participants

A thorough description of the current sample, including exclusion due to low rs-fMRI data quality is reported elsewhere (47; see Supplemental Materials). In short, we retained a final sample of 82 age- and sex-matched adolescents and young adults (n_BPD_ = 40, mean age = 20.53, age range 13-30) with and without clinically heightened BPD symptoms. Detailed demographic information can be found in Table 1.

**Table 1.** Sample Characteristics

| Characteristic | BPD (*n* = 40) | HC (*n* = 42) |
| --- | --- | --- |
| Age (SD) | 20.84 (4.42) | 20.61 (4.16) |
| PAI-BOR (SD) | 47.3 (9.60) | 10.3 (3.99) |
| Ethnicity |  |  |
| Hispanic or Latino | 3 | 2 |
| Not Hispanic or Latino | 36 | 40 |
| Not Provided/Missing | 1 | 0 |
| Race |  |  |
| Caucasian | 31 | 30 |
| African American | 3 | 7 |
| Asian American | 2 | 1 |
| Bi/Multiracial | 2 | 4 |
| Not Provided/Missing | 2 | 0 |
| Average Annual Income |  |  |
| < \$5,000-\$19,999 | 10 | 11 |
| \$20,000-\$34,999 | 9 | 7 |
| \$35,000 - \$59,999 | 8 | 5 |
| \$60,000 - \$99,999 | 5 | 6 |
| \$100,000 + | 3 | 10 |
| Not Provided/Missing | 5 | 3 |
| Sexuality |  |  |
| Heterosexual | 28 | 40 |
| Gay/Lesbian | 1 | 1 |
| Bisexual | 8 | 0 |
| Other | 1 | 1 |
| Not Provided/Missing | 2 | 0 |
| Psychiatric Medication |  |  |
| Any Psychiatric Medication | 18 | 0 |
| SSRI | 15 | 0 |
| SNRI | 2 | 0 |
| Bupropion | 3 | 0 |
| Sedative | 5 | 0 |
| Antipsychotic | 0 | 0 |
| Anticonvulsant | 2 | 0 |

*Note.* Samples were sex- and age-matched. PAI-BOR: Personality Assessment Inventory – Borderline subscale.

### Procedure

Participants completed two semi-structured diagnostic interviews assessing psychopathology and personality disorder symptoms (55,56). Interviews were administered by research assistants trained and supervised by the senior author. Participants in the BPD group met diagnostic criteria for three or more of the DSM-IV-TR BPD symptoms (47,57). Exclusionary criteria included having a first-degree relative diagnosed with Bipolar I disorder or any psychotic disorder and a history of serious head injury or neurological disease. In addition, control participants had no history of psychiatric or substance abuse disorders.

Prior to the RS-fMRI session, participants completed a battery of self-report measures including the Borderline Personality Questionnaire (BPQ; 58). The BPQ is an 80-item self-report measure which measures the various BPD symptom dimensions based on the nine DSM criteria. Internal consistency among these scales was good to excellent in our sample (α_BPQ_ = 0.97, mean subscale α_BPQ_ = 0.86).

### MR Data Acquisition

Data were acquired using a Siemens 3T Tim Trio scanner with a 32-channel head coil at the University of Pittsburgh Medical Center. We collected five minutes of resting-state fMRI data during which subjects were asked to keep their eyes open and relax, but not fall asleep. We used a simultaneous multi-slice echo-planar sequence sensitive to BOLD contrast with scanning parameters: TR = 1.0s, TE = 30ms, FoV = 220 mm, flip angle = 55°, voxel size = 2.3mm isotropic, 5x multiband acceleration. We confirmed that no subjects fell asleep with a self-report questionnaire administered after the scanning protocol.

### RS-fMRI Preprocessing Procedures

RS-fMRI preprocessing was conducted within FSL, NiPy, and AFNI. Structural scans were registered to the MNI152 template (59) using affine and nonlinear transformations conducted in FSL. Functional image preprocessing included simultaneous 4-D motion and slice-timing correction (60), brain extraction, alignment of subject’s functional images to their anatomical scan using a boundary-based registration algorithm (61), and a one-step nonlinear warp to MNI152 space that concatenated functional-to structural, structural-to-MNI152, and fieldmap unwarping transformations. To mitigate motion-related artifacts we used ICA-AROMA (62), a data-driven classification algorithm that identifies and removes spatiotemporal components likely to reflect head movement. RS-fMRI data were not spatially smoothed for analysis (Supplemental Methods;, 63).

### Analytic Approach

#### Whole-brain Functional Connectivity and ALFF Analyses

We performed whole-brain FC analyses (see Supplemental Methods for details) to identify resting-state network signatures of BPD across levels of the network. We first constructed undirected FC matrices among 421 regions/nodes using a custom-built parcellation covering cortex, striatum, thalamus, and amygdala [43, 64–66; see Fig S2]. Given our interest in network segregation and integration, nodes were assigned to either the default mode (DMN), fronto-parietal (FPN), salience (SN), dorsal attention (DAN), sommato-motor (SomMot), visual (Vis), or cortico-limbic network (Limbic), based on the previously validated modular structure of the cortical and striatal parcellations (64,65,67).

To test for group differences in global FC and ALFF, we compared the strength centrality and global ALFF distributions of the two groups using mixed-effects regression (see Supplemental Methods). We calculated graph measures of global FC (modularity, characteristic path length, transitivity, global efficiency, diameter) and fit separate regression models predicting global graph metrics by group membership, age, and their interaction. We then computed nodal FC measures and ALFF for all brain regions (see Supplemental Methods). In a series of nine logistic ridge regression models, we tested which regional connectivity measures were the most potent predictors of group status; these analyses included graph measures for each region as simultaneous predictors (see Supplemental Methods). Nodal measures included strength centrality, seven separate network-specific strength centrality (NSSC) scores, and ALFF. The NSSC measures corresponded to intra- and inter-network FC with specific networks. For nodal logistic ridge regression analyses we set a conservative alpha level of .005.

As detailed below, we found strong evidence for widespread hyperconnectivity of the R daIns in BPD. This result motivated a post-hoc analysis on all edges incident to this region, testing for edges with significant group or group-by-age effects. The focal edge analysis provided a more fine-grained test of *which* edges contributed most to group differences in summary statistics such as nodal centrality. Specifically, in a logistic ridge regression, we regressed group status on incident edge values and their interactions with age.

#### Effective Connectivity Among Target Regions

Initial analyses revealed pivotal group differences in FC/ALFF in R daIns, TPJ, and several (primarily parietal) DAN regions. Subsequent analyses sought to understand how these effects could be understood in terms of EC amongst these regions. We used the recently developed Latent Variable Group Iterative Multiple Model Estimation (LV-GIMME; 44,68,69) algorithm to estimate EC between R TPJ, daIns, and a latent variable capturing shared signal amongst DAN regions (see Supplemental Methods). We additionally allowed for subgroup-specific edges to be estimated based on clinical group status (69). After fitting LV-GIMME to nodal time series, we extracted model coefficients as group-level directed edge values and ran separate linear regressions testing for group and group-by-age effects in the strength of EC.

#### Path Models Linking EC and ALFF

Building on results from EC estimation, we tested whether the relationship between DAN_ALFF_^3^ and daIns_ALFF_/TPJ_ALFF_ were mediated by EC between these regions^4^. For example, while daIns_ALFF_ and DAN_ALFF_ levels showed an inverse pattern of group-level effects, we directly considered whether parametric increases in daIns daIns→DAN_EC_ inversely scales with DAN_ALFF_ (which would demonstrate an “ALFF suppression” effect).

We fit three path models to these variables in Mplus using Bayesian parameter estimation (details in Supplemental Methods; 69–71). In the first two models (Fig. 2a-b), we fit parallel dual-mediation path models in which the relationship between daIns_ALFF_/TPJ_ALFF_ and DAN_ALFF_ was mediated by both daIns/TPJ→DAN_EC_ and daIns/TPJ↔DAN_FC_, respectively (Fig. 2a-b). The inclusion of FC and EC measures tested the specificity and predictive power of directed and undirected connectivity. In a final model (Fig. 2c), we tested whether TPJ-related variables were predicted by daIns→TPJ_EC_. This path model specifically adjudicated whether TPJ_ALFF_ and/or TPJ connectivity were better explained as a downstream effect of daIns hyperactivity/connectivity.

#### Associations with BPD Symptom Domains

Finally, we tested which BPD symptom dimensions were uniquely associated with ALFF impairment in DAN. We fit a series of linear models using each BPQ subscale: 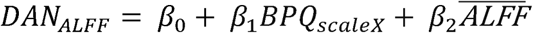. However, we were more interested in the unique relevance of specific BPQ subscales given high associations among symptom domains (*r*’s > .7). To test conditional associations of BPQ subscales with DAN ALFF we fit a single multiple regression predicting levels of DAN ALFF by all subscales of the BPQ and average ALFF levels per subject^5^.

## Results

### Whole-brain functional connectivity and ALFF

We found no evidence of significant group or group-by-age differences in global network characteristics (all *p*’s > .05; Table S2, Figs S4-5). Nodal analyses, however, revealed an array of findings across intrinsic networks (Table S3, Fig S6). The primary goal of our nodal analyses was to determine group and group x age differences in FC/ALFF across regions.

We identified three robust patterns in intrinsic network structure that discriminated the BPD group from the control group (Table S3, Fig S6). First, R dorsal aIns (daIns) showed strong, widespread hyperconnectivity to six of seven^6^ intrinsic networks. Most notably, R daIns↔DAN_FC_ was substantially stronger in the BPD group compared to controls (*t* = 4.55, *p* < .0001), controlling for all nodes’ NSSC_DAN_ scores. In addition, daIns_ALFF_ was significantly higher in the BPD group (*t* = 3.76, *p* < .001). Second, we found a pattern of hypoconnectivity in R TPJ, as indicated by lower strength centrality in the BPD group (*t* = −2.99, *p* < .005). Third, we found lower ALFF in the BPD group across parietal nodes in the DAN (all *p*’s < .005).

We conducted a post-hoc ridge regression of all edges connected to daIns^7^. This analysis revealed hyperconnectivity between R daIns and 11 DAN regions (Table S4, all *p*’s < .005), suggesting a broad pattern of hyperconnectivity between daIns and DAN. Parietal DAN regions with lower ALFF overlapped considerably with nodes exhibiting heightened connectivity with daIns (Table S6, Fig S7).

### Effective Connectivity Amongst Target Regions

Whole-brain FC/ALFF effects suggested a robust pattern of heightened daIns↔DAN_FC_ coupled with impaired DAN_ALFF_ in the BPD group. However, these analyses cannot reveal the direction of information flow between daIns and DAN. Given insula’s status as a major “outflow hub” (18), we interrogated if heightened daIns↔DAN_FC_ could be better understood in terms daIns→DAN_EC_, rather than DAN→daIns_EC_. To test this hypothesis, we estimated EC among R daIns, TPJ, and 15 selected DAN regions using LV-GIMME (Fig 1). DAN regions were selected based on the unison of regions with lower ALFF *or* heightened FC to daIns (details in Supplemental Materials).

**Figure 1.**
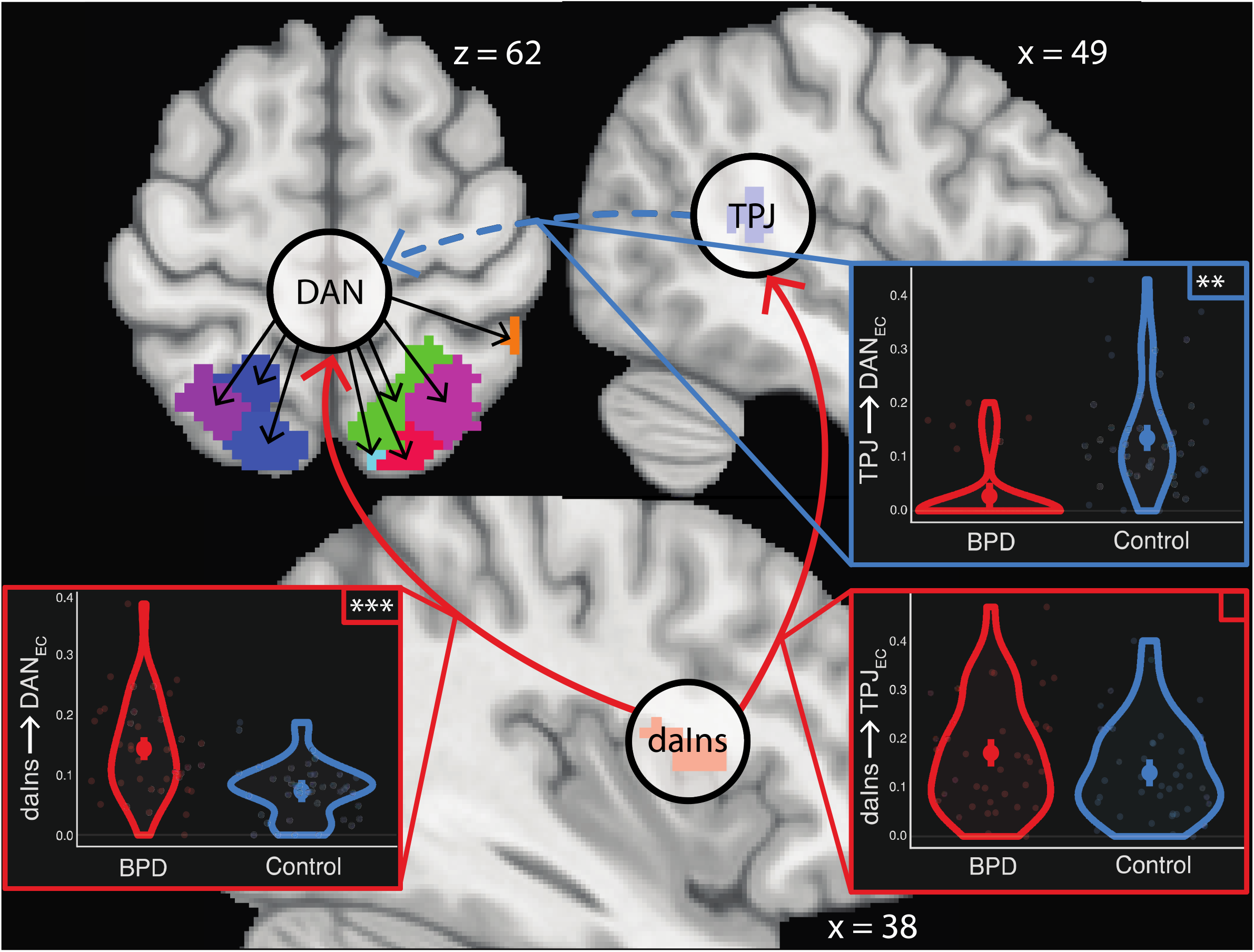
Effective connectivity results. *Note*. Circles denote BOLD signals fit within LV-GIMME framework. Note that SN nodes (daIns, TPJ) are fit to the nodal timeseries whereas DAN was fit as a network-level latent variable given the strong pattern of shared signal across DAN regions (see Table S6). Red solid arrows denote group-level edges that were significantly higher in the BPD group, where the dashed blue arrow denotes a HC-specific edge from TPJ to DAN. Violin/density plots are overlain on extracted model betas for each edge (e.g. model-predicted edge values) and bolded points denote the expected marginal mean of each edge for each group, averaged over mean FC value (***p < .001, *p < .05; details in Table S7).

To capture the shared signal among parietal regions in DAN, we fit a latent variable model to their timeseries with LV-GIMME. Results demonstrated that daIns had a directed influence on both DAN and R TPJ (Fig 1). Further, LV-GIMME detected TPJ→DAN_EC_ in HCs that was not present in the BPD group. Linear models indicated that daIns→DAN_EC_ and daIns→TPJ_EC_ were significantly higher in borderline participants (t_DAN_ = 5.27, p_DAN_ < .001; t_TPJ_ = 2.04, p_TPJ_ = .04; Table S7).

### Path Models Linking EC and ALFF

Having established group-level increases in daIns→DAN_EC_ and daIns→TPJ_EC_ and widespread lower DAN_ALFF_ in the BPD group, we integrated these findings with a set of multivariate path analyses. In our daIns→DAN path model (Table S8, Fig 2a) we found modest evidence that DAN_ALFF_ was negatively associated with daIns_ALFF_ (*p* = 0.05) and daIns→DAN_EC_ (p = 0.06), but not daIns↔DAN_FC_ (p = 0.83), controlling for subject-level average FC and ALFF. Further, daIns→DAN_EC_ was positively associated with daIns_ALFF_ (p < 0.001). We found limited support that daIns→DAN_EC_ mediates the relationship between daIns_ALFF_ and DAN_ALFF_ (*p* = 0.07). In a parallel model (TPJ→DAN, Fig 2b), we tested the associations between TPJ and DAN connectivity and ALFF. We found that TPJ→DAN_EC_ was associated with higher DAN_ALFF_ (*p* = 0.02). DAN_ALFF_ was not associated with TPJ_ALFF_ (p = 0.38) or TPJ↔DAN_FC_ (p = 0.73). Further, TPJ_ALFF_ was associated with higher TPJ→DAN_EC_ (p < 0.001) and TPJ↔DAN_FC_ (p < 0.001). The association between TPJ_ALFF_ and DAN_ALFF_ was fully mediated by TPJ→DAN_EC_ (p = 0.02).

**Figure 2.**
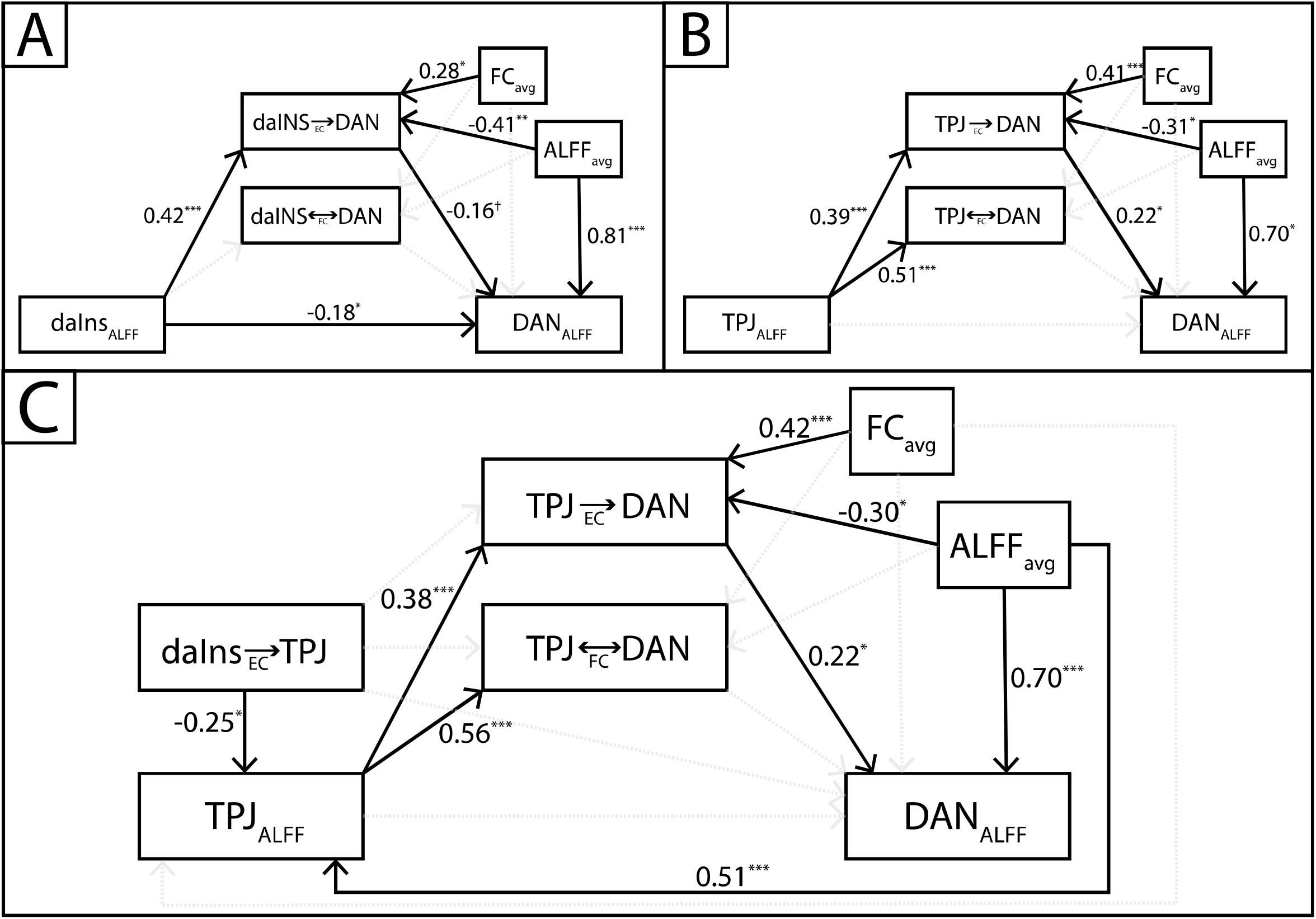
Graphical depiction of path models linking daIns, TPJ, and DAN signals. *Note*. Visual depiction of path models, fit with Bayesian parameter estimation in Mplus. Path values denote standardized estimates. In top left and right panels, we depict parallel dual-mediator path models that tested the ability of SN regions (daIns on left and TPJ on right) FC *and* EC to DAN to mediate the relationship between daIns_ALFF_/TPJ_ALFF_ and DAN_ALFF_, with both models finding support that EC from both SN regions mediate this relationship and outcompete FC measures in explaining variance in DAN_ALFF_. In the bottom panel, our combined model builds on the TPJ → DAN model, by showing that EC from daIns to TPJ suppresses TPJ_ALFF_ leading to downstream consequences on TPJ to DAN EC. Details on all three models are reported in Table 3 (combined) and Tables S8-9 (initial parallel mediation models).

In our daIns→TPJ→DAN model (Table 2, Fig 2c), we built upon the TPJ→DAN model^8^ by demonstrating that TPJ→DAN_EC_ depends on daIns→TPJ_EC_. While daIns→TPJ_EC_ did not directly predict TPJ→DAN_EC_ (p = 0.74), or TPJ↔DAN_FC_ (p = 0.17), it significantly predicted weaker TPJ_ALFF_ (p = 0.02). In turn, TPJ_ALFF_ fully mediated the relationship between daIns→TPJ_EC_ and both TPJ→DAN_EC_ (p = 0.02) and TPJ↔DAN_FC_ (p = 0.02).

**Table 2.**
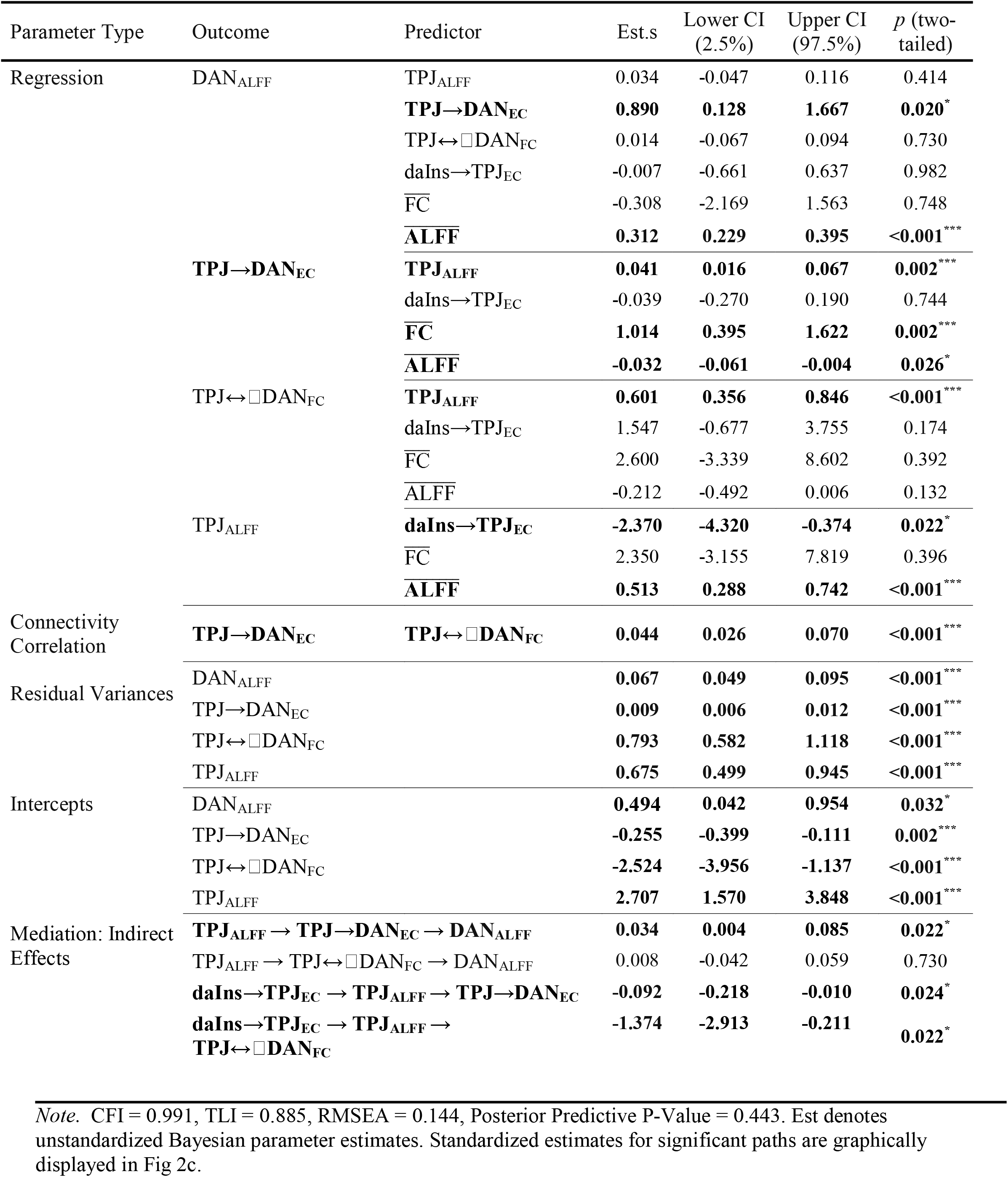
Parameter Table: daIns→TPJ→DAN (Combined) Model

### Associations of connectivity measures with BPD Symptom Domains

EC and path analyses demonstrated that lower DAN_ALFF_ in the BPD group is a downstream effect of differential daIns→DAN_EC_ and TPJ→DAN_EC_. Building on this, we sought to test which borderline symptom domains were most associated with DAN_ALFF_ impairment. In unconditional models, nearly every BPQ subscale was negatively associated with DAN_ALFF_ (Table 3). However, in a single conditional model, the negative association between DAN_ALFF_ and the affective instability subscale remained significant, while the other subscales were not (all *p*’s > .19, Table 3).

**Table 3.** DAN_ALFF_ is negatively associated with higher affective instability

| Predictor | $t_{\text{unconditional}}$ | $p_{\text{unconditional}}$ | $t_{\text{conditional}}$ | $p_{\text{conditional}}$ | $\overline{R^2}_{\text{dom}}$ |
| --- | --- | --- | --- | --- | --- |
| Impulsivity<sub>BPQ</sub> | -2.64 | 0.010** | -1.07 | 0.291 | 0.009 |
| **Affective Instability<sub>BPQ</sub>** | **-3.97** | **0.002\*\*** | **-2.87** | **0.005\*\*** | **0.035** |
| Abandonment<sub>BPQ</sub> | -2.38 | 0.020* | -0.04 | 0.970 | 0.009 |
| Relationships<sub>BPQ</sub> | -2.68 | 0.009** | -0.24 | 0.815 | 0.017 |
| Self-Image<sub>BPQ</sub> | -2.13 | 0.036* | 0.29 | 0.771 | 0.020 |
| Suicide/Self-Mutilation<sub>BPQ</sub> | -2.04 | 0.045* | 1.32 | 0.192 | 0.012 |
| Emptiness<sub>BPQ</sub> | -2.08 | 0.041* | -0.08 | 0.936 | 0.006 |
| Intense Anger<sub>BPQ</sub> | -1.80 | 0.075<sup>†</sup> | 0.68 | 0.500 | 0.004 |
| Quasi-Psychotic States<sub>BPQ</sub> | -1.15 | 0.256 | 0.63 | 0.532 | 0.010 |
| <u>ALFF</u> | 11.75 | <0.001*** | 11.30 | <0.001*** | 0.590 |

*Note.* Predictor column reflects BPQ subscales that were used to predict DAN<sub>ALFF</sub>. ALFF represents a nuisance covariate calculated as a subject-specific mean of regional ALFF values. $t_{\text{unconditional}}$ and $p_{\text{unconditional}}$ reflect test statistic and p-value for separately fit linear models for each BPQ subscale, where $t_{\text{conditional}}$ and $p_{\text{conditional}}$ reflect test statistic and p-value for parameters of a joint multiple regression, where subscales compete to explain variance in DAN<sub>ALFF</sub> (see *Associations with BPD Symptom Domains* in Methods). $\overline{R^2}_{\text{dom}}$ represents the average change in model $R^2$ when entering this symptom scale into nested sub-models in a dominance analysis signaling the relative importance of heightened affective instability in predicting lower DAN<sub>ALFF</sub> compared to other BPD symptom domains.

## Discussion

BPD symptoms often emerge during adolescence (2), a period associated with major neurodevelopmental changes. In a sample of adolescents and young adults with BPD symptoms, we found that right dorsal anterior insula (R daIns) was hyperactive (ALFF) and hyperconnected (FC and EC) to DAN relative to matched controls. We also found hypoactivity across parietal regions in DAN in BPD participants. Path analyses revealed that heightened daIns_ALFF_ (53) bolstered daIns→DAN_EC_. Importantly, heightened daIns→DAN_EC_ negatively predicted DAN_ALFF_ over and above daIns↔DAN_FC_. Our findings suggest that blunted resting-state activity in DAN may be a useful biomarker of BPD that reflects differential inputs from SN regions. Comparing across BPD symptom domains, diminished DAN_ALFF_ was most strongly associated with affective instability. Although we allowed age to moderate these effects, our primary results were consistent across the age range of the sample (13–30). Thus, differences in daIns connectivity and DAN_ALFF_ may be a characteristic of BPD more generally.

We found that lower DAN_ALFF_ was differentially predicted by EC from two key SN nodes: whereas daIns→DAN_EC_ suppressed DAN_ALFF_, TPJ→DAN_EC_ enhanced DAN_ALFF_. Although DAN_ALFF_ was bolstered by TPJ→DAN_EC_, TPJ_ALFF_ was itself suppressed by daIns→TPJ_EC_ in the BPD group. Thus, while daIns→DAN_EC_ and TPJ→DAN_EC_ play competing roles in modulating DAN_ALFF_, our analyses suggest the primacy of daIns in modulating activity in both DAN and TPJ. These results provide compelling evidence for aberrant integration of SN and DAN in BPD. A more nuanced interpretation is that abnormal intra-network communication within the SN (stemming from increased daIns→TPJ_EC_) leads to an imbalance in the strength of inputs from anterior (insula) vs posterior (TPJ) nodes of the SN to DAN.

aIns is a core node of SN that supports cognitive-emotional processing via attentional shifts that accord with task-related goals by assigning salience to behaviorally relevant cues (15,17). Its rapid integration of visceral/homeostatic, emotional, social, cognitive, and sensory signals place aIns in a unique position to control the ongoing assignment of resources to networks responsible for the processing of exogenous cues or internal self-referential information (15,18,20). In previous studies daIns showed the strongest pattern of co-activation with higher-order cognitive networks (FPN, DAN) and was more strongly associated with switching tasks, whereas ventral anterior and posterior insula co-activate with affective and sensorimotor processing networks, respectively (21). This evidence aligns well with our finding of greater daIns↔DAN_FC_ in the BPD group, suggesting that hyperactivity in this region has the greatest ability to influence cognitive/executive functions.

We also found that R TPJ, a node of the posterior SN, showed widespread, though less pronounced, hypoconnectivity across networks. Prior fMRI studies of BPD have found that R TPJ is hypoactivated across tasks measuring self/other differentiation, perspective-taking, and social feedback processing (29,74–76). These findings align well with R TPJ’s hypothesized role in theory of mind and mentalization (77) and more recent claims that TPJ is crucial in constructing social contexts (78) and representing social agents and their intentions (79). Hypoconnectivity of R TPJ in our data may reflect difficulties in perspective-taking and mentalizing associated with adolescent (80–82) and adult (83) BPD. However, TPJ is also linked to the more universal task of contextual updating and adjusting top-down expectations (84), providing an intriguing clue for a domain-general functional impairment in BPD that would have direct implications for social-cognitive functioning. The hyper-vs hypo-connectivity dissociation within SN (daIns versus TPJ) helps to reconcile previous conflicting findings of SN connectivity in BPD. One ICA study found increased FC of canonical/anterior SN, including insula and dACC (34), where another study found hypoconnectivity in a more widespread “social salience network” that included canonical SN nodes *and* TPJ (33). While R daIns and TPJ appear to form a functional circuit (i.e. SN), intra-SN daIns→TPJ_EC_ impairs TPJ_ALFF_, leading to downstream impairment of inter-network TPJ→DAN_EC_ in BPD participants.

DAN is involved in the control of goal-oriented/top-down selective attention (85,86) and plays a central role in prioritizing sensory inputs for further processing, including saccades towards high priority stimuli (87). Extending this notion, parietal DAN nodes construct a “priority map” of visual input by integrating top-down signals relevant to goals and expectations with a compressed representation of sensory features. In turn, priority maps bias neural activity in primary sensory networks and govern information-seeking behaviors, particularly saccades (88–90). Lower DAN_ALFF_ in BPD points to a diminished capacity to integrate higher-order goal representations to guide attention. Thus, attentional orientation towards stimuli that are associated with abstract goals may be overpowered by daIns-generated switch signals to brief stimuli with high associative salience. This account is partially consistent with the triple network model, which posits that altered salience processing in aIns underlies key dysfunctional interactions between FPN, SN, and DMN. However, we found stronger evidence of aberrant SN↔DAN_FC_, providing an intriguing extension of the triple network model in borderline personality (23). Our data supports a slightly different view that aberrant SN functioning has greater direct implications on selective or transient attentional control implemented by DAN (91), which shows anatomical and functional dissociations from FPN (39,92).

We propose that heightened daIns→DAN_EC_ in BPD may reflect a vulnerability to “hijacking” of goal-directed, transient attention by the aIns. Heightened activity and connectivity of daIns may increase attentional switches among competing priorities or immediate needs in young people with BPD symptoms. This attentional hijacking hypothesis, if corroborated and extended in subsequent research, could provide a circuit-based account of attentional biases in BPD, characterized by faster saccades to and longer fixations on the eyes of emotional faces (93–95). While previous FC studies of SN-DAN interactions remain equivocal on the directionality of influence between these networks, our LV-GIMME analysis specifically tested the hypothesis that SN nodes primarily act on DAN. Our analysis suggests that in individuals with BPD symptoms, TPJ_ALFF_ and DAN_ALFF_ both fall prey to hijacking by daIns. A high-level conclusion from our analyses is that heightened switch-signaling from daIns impairs DAN_ALFF_ both directly (daIns→DAN_EC_) and indirectly vis-à-vis blunted TPJ→DAN_EC_. Moreover, weaker DAN_ALFF_ was particularly associated with heightened affective instability in our study, suggesting that disrupted intrinsic activity in DAN enhances vulnerability to negative emotion (96).

To our knowledge this is the first whole-brain rsFC study in a sample of adolescents and young adults with BPD symptoms, allowing for an initial description of intrinsic connectivity during a period of heightened vulnerability. Second, our study combined well-validated parcellations of the cortex, striatum, amygdala, and thalamus, allowing for a fine-grained analysis of whole-brain FC. Third, our network neuroscience approach leveraged NSSC scores, allowing for focal tests of intra- and inter-network connectivity that overcome the limitations of broader between-network connectivity measures such as participation coefficient (97). Fourth, we analyzed ALFF to uncover differential contributions of intrinsic connectivity and intrinsic activity to the functional network profile of BPD. Fifth, our LV-GIMME analysis reflects a state-of-the-art approach to measuring the directionality of connectivity when there are shared signals (here, nodes within DAN) in a network (98).

While our results suggest a novel “attentional hijacking” hypothesis of BPD, in the absence of goal-directed attentional control tasks, this hypothesis remains speculative. Future studies can directly test this hypothesis in task-based studies that manipulate exogenous and/or internal shifts of attention. Our primary findings emphasize group differences that were consistent across individuals between ages 13 and 30, and longitudinal assessments with clinical comparison groups are necessary to directly investigate within-person connectivity *changes* over adolescence. Likewise, replicating these findings in a sample of older adults with BPD symptoms would clarify whether daIns connectivity plays a major role in DAN intrinsic activity across the lifespan.

In summary, we found that intrinsic hypoactivity of DAN plays a pivotal role in the expression of borderline personality symptoms, particularly affective instability, in young people. We suggest that this pattern may represent an insula-driven attentional hijacking process whereby daIns interferes with goal-directed attentional control in DAN both directly and indirectly via impaired TPJ connectivity. This account builds on evidence that the aIns implements attentional switches in response to dynamically changing homeostatic needs on short timescales (13). If hyperconnectivity of the daIns with DAN regions supports rapid attentional shifts, our findings may help explain the well-documented sensitivity to brief emotional cues that form a core feature of borderline personality (99,100). We hope that future studies extend these findings to refine a circuit-based account of borderline symptoms in adolescents that can inform early intervention treatments in this high-risk, yet understudied, population.

## Acknowledgments

This work was funded by the National Institutes of Mental Health (K01 MH097091 and R01 MH119399 to MNH). The funding agency had no role in the design and conduct of the study; the collection, management, analysis, and interpretation of the data; the preparation, review, and approval of the manuscript; or the decision to submit the manuscript for publication.

## Footnotes

1 For a summary of findings from all published rsFC studies of adults with BPD we refer the interested reader to Table S1.

2 Throughout the rest of this paper, we use ALFF and “resting-state activity” interchangeably, though we encourage readers to keep in mind that this perhaps a rough equivalence.

3 Given that ALFF effects in DAN were spread across a wide swath of parietal and frontal eye field regions (Fig S6, Table S3), we were more interested in a shared network-level score for ALFF. We fit a single factor EFA to ALFF scores amongst DAN regions with significant group differences and extracted factor scores to be used at the dependent variable in subsequent analyses. Details including bivariate correlations amongst DAN nodes and factor loadings are detailed in Supplemental Materials.

4 For ease of communication, for the rest of the manuscript we denote effective connectivity between two regions with an arrow and “EC” subscript, and functional connectivity between two regions/networks with a dash and “FC” subscript (e.g. EC from daIns to DAN denoted as daIns→DAN_EC_, where undirected FC is denoted daIns↔DAN_FC_)

5 This allowed us to test which specific BPD symptom domains explain significant variability in DAN ALFF, while accounting for their correlation with one another. To allay concerns about multicollinearity and the interpretability of partial betas, we report a dominance analysis in order to test for the incremental predictive ability of BPQ subscales (73). Results confirm that the regressor with the largest partial beta in our joint model induces the greatest average increase the model R^2^ across nested sub-models (Table 3).

6 The only network that did not appear to show significantly heightened FC to daIns in the BPD group was the Limbic network. We note that this network showed had a remarkably weaker edge strength distribution compared to the other six networks (visually depicted in Figure S3). This network is comprised primarily of inferior temporal and orbitofrontal regions and thus the regional timeseries are likely much noisier do to susceptibility artifacts in fMRI data.

7 After our mild consensus thresholding procedure (see Supplemental Materials), 26 edges connected to daIns were removed, leaving 394 remaining edges to investigate in our edge analysis.

8 Before moving to interpreting newly introduced daIns paths we confirmed that paths that were present in both models 2 and 3 were nearly identical in terms of parameter values and significance values (Fig 2, Tables S8-9).

