## Supplemental Materials for "Right anterior insula effective connectivity impairs intrinsic BOLD fluctuations in dorsal attention network in adolescents and young adults with borderline personality symptoms"

**Supplemental Methods**

**Participants**

Participants were 46 adolescents and emerging adults with BPD symptoms recruited from community and outpatient settings, as well as 44 sex- and age-matched healthy controls. All participants were screened using the Personality Assessment Inventory-Borderline scale (1) with BPD participants screening ≥ 30 and controls screening < 17. The average age was 20.53 years (range 13-30 years); 59 participants were female and 31 were male. Eight participants (*n* BPD = 6) were excluded from our analyses for having poor fMRI data quality due to excessive motion during the scan or failure to pass residual correlation checks (see fMRI QA and exclusion). See Table 1 for a complete demographic characterization of the final sample.

**fMRI QA and Exclusion**

For all participants, we calculated volume-to-volume framewise displacement (2). We excluded subjects with FD > 0.5mm in at least 20% of the volumes, or any FD > 10mm. This led to the removal of six participants, four of whom were in the BPD group. We removed one additional subject from the BPD group whose data did not pass serial residual correlation checks after pre-whitening our data. We further removed one participant from the BPD group who passed our head motion criteria but whose functional connectivity matrix was remarkably different from the group average (see Fig S1), leaving a final sample of 82 participants.

**RS-fMRI preprocessing procedures**

Although many resting-state fMRI studies apply spatial smoothing, we analyzed unsmoothed data because 1) our whole-brain parcellation encompasses the entire brain volume, leading to the spatial proximity of many nodes, and 2) a recent report documenting that spatial smoothing alters graph theory-based measured of connectivity non-uniformly across the brain (3). Although our connectivity analyses were conducted on unsmoothed data, ICA-AROMA was conducted on data that were spatially smoothed with a 5mm FWHM gaussian kernel (FSL *susan*), consistent with recommended guidelines. Spatial smoothing increases SNR in BOLD data, allowing for an increased ability to detect structured artifacts that should be removed from the signal (such as components related to subject movement; 1). Based on the results of ICA-AROMA, we regressed motion-related components out of the unsmoothed data using *fsl* *regfilt* (i.e. “non-aggressive” denoising). AROMA’s automated component selection approach has recently been shown to be superior to other competing procedures in removing motion artefacts while preserving the signal of interest, and it largely eliminates distance-dependent motion-FC correlation effects (4–6).

**Nodal parcellation**

Prior to conducting graph analysis, we parceled voxels into 421 functional regions (nodes) using a custom-built atlas (Fig S2) by combining state-of-the-art functional parcellations of the cortex (7), striatum (8), thalamus (9), and amygdala.(10,11). We parcellated voxels by assigning atlas values (node numbers), via a combined parcellation mask, with the goal of eventually averaging voxel timeseries within a given node to obtain an average timeseries per node and subject. Before computing nodal timeseries, we calculated subject-level masks that reflected the proportion of voxels in each ROI that contained unreliable signal, as indicated by voxelwise standard deviation equal to zero and/or all values equal to zero. Visual inspection of the subject-level masks indicated that problematic voxels were located predominantly in inferior temporal regions and to a lesser extent, orbitofrontal regions, reflecting signal loss due to susceptibility (see Fig S2). We then merged all binary masks into a group-level mask with voxel-wise values equal to the proportion of subjects with reliable signal in the voxel. In order to ensure that ROI time series reflected the same voxels across participants, we removed all voxels from the parcellation in which less than 95% of subjects had reliable signal. This procedure removed 600 voxels —approximately .7% of the total voxels — from our parcellation.

**Prewhitening** **nodal timeseries and adjacency (undirected FC) matrix construction**

After performing motion removal procedures and constructing our combined parcellation, we pre-whitened our nodal time series using a series of increasingly complex ARMA models prior to calculating functional connectivity to ensure that cross-correlation estimates were not biased by the temporal (i.e. autocorrelated) structure of BOLD data. This decision was based on the concern that failing to remove autoregressive components of fMRI time-series violates a key assumption of the general linear model (specifically, residuals must be *i.i.d*. and normally distributed; 8). Furthermore, estimates of cross-correlation can be misestimated when time series have similar autoregressive properties that do not reflect true interregional connectivity. To overcome this concern, auto-regressive models such as Auto-Regressive Moving Average (ARMA(p,q); 9) models have received increasing attention in the fMRI literature in recent years, where high amounts of serial correlation are inherent in the data structure. Prior to running ARMA models, we computed aggregated nodal time series by taking the average of all time series for voxels included in a given node, excluding those voxels that were missing or had no variance (see above).

ARMA models represent the temporal dependence of observations in a time series, allowing one to remove the autoregressive components of the signal to achieve a “white” error time series. ARMA models were fit to each average nodal time series using the *Arima* function included in the *forecast* package in R (14) such that

$$\eta_{t}=c+\varepsilon_{t}+\sum_{i=1}^{p} \varphi_{i}\eta_{t-i}+ \sum_{i=1}^{q} \theta_{i}\varepsilon_{t-i}$$

where $\varphi_{i}\ldots\varphi_{p}$ denote freely estimated AR coefficients that quantify the degree of autocorrelation between the current realization ($\eta_{t}$) and previous realizations, and $\theta_{i}\ldots\theta_{p}$ denote freely estimated MA coefficients that quantify the degree of dependence of the current innovation ($\varepsilon_{t}$) on prior innovations. The residual error term, also called the innovation term in an AR model, is assumed to be normally distributed (i.e. “white”) noise

$\varepsilon_{t}\sim iid N(0,\sigma^{2})$.

The *unique* coefficients of a given ARMA model for a node and subject act as a filter on the time series that whitens the residuals. However, the important quantity of interest in a graph theoretical analysis is the cross-correlation between every pair of nodal time series within a given subject, which are then used as cells in the adjacency matrix. Thus, to compute functional connectivity between regions, we stored the ARMA coefficients (sometimes referred to as the transfer function) for one node and used these to filter both time series for a given subject. We used the Pearson product-moment correlation coefficient to quantify the functional connectivity between nodal time series that had been passed through the same ARMA coefficients:

$$r_{\phi\left( y \right),\phi(x)}= \frac{1}{T-1} \sum_{t=1}^{T} \frac{\phi\left( y_{t} \right)-\bar{\phi(y)}}{s_{\phi(y)}}\cdot\frac{\phi\left( x_{t} \right)- \bar{\phi(x)}}{s_{\phi(x)}}$$

where $\phi\left( y \right)$ and $\phi\left( x \right)$ denote the time series for two nodes that have both been filtered by the fitted ARMA coefficients for *y*.

We fit a series of increasing complex ARMA models until the number of subject-wide “non-white” residuals fell below 5% of voxels for all subjects (i.e., the false positive rate on the test). Time series were deemed “non-white” on the basis of the Breusch-Godfrey test, which tested null hypotheses of serial correlation our nodal time series, which was computed up to six lags prior to the current realization (i.e., six seconds in the past). Through this procedure we retained our results from an ARMA(4,2) model as the edges of subject-level graphs for further analysis. We excluded one subject from all analyses who’s residual time series remained non-white after ARMA(4,2) pre-whitening, whereas in all other subjects our prewhitening procedures were successful.

Finally, after identifying the correct model order, we computed the Pearson correlations (described above) among the pre-whitened timeseries (residuals of ARMA(4,2)) to yield a 421x421 adjacency matrix for each subject representing FC amongst our 421 nodes. In order to remove unreliable edges from these matrices, we applied a minimal consensus thresholding procedure (15). Specifically, we removed edges from all subjects that did not have a weight of *r* = .1 or higher in 25% or more of subjects.

**Intrinsic network (module/sub-network) assignment**

In order to compare levels of connectivity within and between intrinsic networks of interest, we assigned nodes to one of seven canonical intrinsic networks based on nodal assignments during validation of the cortical parcellation (7,16). Nodes were assigned to either the default mode (DMN), fronto-parietal (FPN), salience/ventral attention (SN/VAN), dorsal attention (DAN), sommato-motor (SomMot), visual (Vis), or cortico-limbic network (Limbic). To represent established cortical-striato-thalamic loops (17), we assigned striatal nodes to intrinsic networks based on their previously reported intrinsic connectivity profiles (8). Subdivisions of the thalamus were assigned to FPN, SomMot, DAN, and DMN based on previously reported prominent white matter projections (9). All amygdala ROIs were assigned to the Limbic network. In order to informally check if within-network FC was indeed greater than between-network FC, we visually inspected within and between network edge distributions and confirmed that this was the case (Fig S3), lending support to the validity of our network partition.

**Analytic approach**

***Whole-brain functional connectivity and ALFF analyses: metric computation***

After computing adjacency matrices, building on prior suggestions (10) we sought to quantify both global and nodal characteristics of subject-specific graphs in order to assess if group-level differences in graph structure are operative at a whole-brain or region/node level. We first computed metrics assessing global and nodal connectivity as well as the amplitude of low-frequency fluctuations (ALFF).

**Global graph characteristics.** At the global level, we tested for group differences and group x age interactions in the strength centrality distributions of the entire network (Fig S4). Presentation of the degree distribution (or its weighted variant strength) was recently proposed as a fundamental property of a graph (10) and can reveal high-level information about global connectivity patterns between groups. Likewise, given our interest in studying ALFF across regions and networks we also tested similar group and group x age effects in global group differences in ALFF.

We further examined several global graph metrics in order to examine the possibility of global graph differences between groups. We calculated weighted modularity, characteristic path length, transitivity, global efficiency, and the weighted diameter of individual graphs (for formulas of weighted variants of these global graph metrics see 18). All computation of global graph metrics was conducted using the R package *igraph* (19).

**Nodal centrality.** Our primary interest in our undirected FC analysis was to identify nodes with markedly different connectivity profiles between groups. Thus, for each subject, at the nodal level we calculated strength centrality, or the sum of all edge weights incident to a node of interest:

$$k_{i}^{w}= \sum_{j \in G} w_{i,j}$$

such that $k_{i}^{w}$ is the weighted degree (strength) centrality for node *i*, and $w_{i,j}$ is the weight of the connection between node *i* and each node *j* for all nodes in the graph (*G*).

In order to focally test connections within and between the seven networks of interest, we also partitioned strength centrality for each node into seven network-specific estimates (one per network):

$$k_{i, N}^{w}= \sum_{j \in N} w_{i,j}.$$

Here, *N* corresponds to the set of all nodes in one of the seven intrinsic networks. Since the seven networks each contain a different number of nodes, we normalized $k_{i, N}^{w}$:

${NSSC}_{i,N}^{w} = \frac{k_{i, N}^{w}- \bar{k_{N}^{w}}}{\sigma_{k_{N}^{w}}}$.

Thus, for $i\in N$, ${NSSC}_{i, N}^{w}$ corresponds to node $i$’s within-module degree *z*-score (20). Otherwise, for $i\notin N{, NSSC}_{i, N}^{w}$corresponds to the normalized inter-network connectivity between node *i* and all nodes in$N$, providing a more specific measure of *which* intrinsic network a node is connected to. Inter-module connectivity is usually assessed via the participation coefficient (20), which provides less specific information about *where* the node’s inter-module connectivity is likely located. Thus, our decision to partition strength centrality into seven distinct network-specific strength scores allows for focal tests of which network a given node may be hyper- or hypo-connected to.

**Nodal amplitude of low frequency fluctuations.** We were also interested in the degree of convergence between connectivity-based measures and the power of low frequency BOLD oscillations (21). This is quantified as the amplitude of low frequency fluctuations (ALFF; 50) and is roughly considered a resting-state analog of RS-fMRI “activity”, due to prior studies linking ALFF to increased regional metabolic demands in RS-fMRI (23). ALFF was calculated for every subject and node as the average power spectrum amongst low-frequency signals (0.01-0.1 Hz; ,21,24) ^[1]^ via:

$$ALFF= \frac{1}{L}\sum_{0.01Hz}^{0.1Hz} Power$$

where the power of a given frequency is obtained via a Fast Fourier Transformation and L denotes the number of Fourier coefficients, thus averaging the summed amplitudes across the selected frequency range (22). ALFF computation was conducted using the *alffmap* function in the *ANTSR* package in R .

***Whole-brain functional connectivity and ALFF analyses: Statistical Analysis***

**Global graph properties.** We first tested for group differences and group x age interaction in the strength centrality and ALFF distributions. Given multiple measures per subject (421 nodal strength estimates) we fit a mixed effects regression (25) predicting strength centrality scores from a combination of between-subjects predictors: binary group, age, and their interaction and fit a random intercept per subject:

$$L1: k^{w}= \beta_{0j}+\beta_{1}Group+\beta_{2}Age+ \beta_{3}Group x Age+ r_{ij}$$

$${L2: \beta}_{0j}= \gamma_{00}+ u_{0j}$$

where $\beta_{0j}$ reflects a subject-specific linear combination of a group-level ($\gamma_{00})$ and subject-specific ($u_{0j})$intercept terms. We replicated the same analysis with the ALFF distribution across nodes, testing the possibility of group or group x age differences in the overall power of low frequency BOLD oscillations. We fit separate multiple regression models with the other global graph characteristics as the DV (five total) using the same fixed effects formula as above (without subject-specific intercepts given one summary score per subject).

**Regularized logistic regression of nodal metrics.** Our primary goal in our whole-brain analysis was to uncover which nodes and nodal metrics (strength, network-specific strengths, and ALFF: nine metrics total) best described BPD-related differences in functional network organization. Given the large number of nodes in our parcellation, we sought to avoid running multiple separate models, which would have a high risk of false positive findings. Moreover, univariate analyses of node centrality would not identify which subset of nodes *jointly* discriminate network differences as a function of BPD and age.

We elected to use a regularized regression approach to overcome the p ≫ n problem in our model (hundreds of parameters, 82 observations), and handle high levels of collinearity in our data (26). Ridge regression shrinks model coefficients towards zero by penalizing the summed parameter estimates. This is achieved by augmenting the standard OLS loss function with an L2 penalty such that

$$L_{ridge}\left( \hat{\beta} \right)= \underset{L_{OLS}}{\underbrace{\sum_{i=1}^{n} {(y_{i}- x_{i}^{'}\hat{\beta})}^{2}}}+\underset{L2 penalty}{\underbrace{\lambda\sum_{j=1}^{m} \hat{\beta}_{j}^{2}}}$$

where λ is a penalty parameter corresponding to the level of shrinkage on the standard OLS regression parameter estimate. The shrinkage parameter λ for each model was chosen using an automated selection algorithm implemented in the *ridge* R package (27,28) and are reported in Table S5.

We estimated nine (one for each nodal metric) logistic ridge regression analyses in which we predicted group status (BPD vs. HC) as a function of nodal metrics for all nodes as well as age-by-metric interactions:

$$logit\left( BPD \right)= \beta_{0}+\beta_{1}Age+ +\beta_{2}{Age}^{2}+ \beta_{3}{FC}_{mean}+ \beta_{4}{NSSC}_{1, DAN}^{w}+ \beta_{5}{NSSC}_{1, DAN}^{w}\times Age+\ldots+\beta_{p-2}{NSSC}_{421, DAN}^{w}+\beta_{p-1}{NSSC}_{421, DAN}^{w} \times Age+\beta_{p}{NSSC}_{421, DAN}^{w} \times{Age}^{2}+e$$

where $p$ is the number of parameters in the model and ${NSSC}_{1, DAN}^{w}$ is a vector of standardized DAN-strength scores for node 1 (denoting the normalized sum of edges connecting node 1 and all nodes in DAN). In this example, all nodes’ connectivity with DAN are fit jointly with ${NSSC}_{i, DAN}^{w}$ of all other nodes, thus partial regression coefficients reflect the unique variability of DAN edges incident to a node to distinguish groups, marginalized over all other DAN edges. In this example, the additional Age^2^ term for ${NSSC}_{421, DAN}^{w}$ denotes that a quadratic age model fit better than linear or inverse variants according to the Vuong test (29). Mean FC value, annual income and average FD per subject were included as nuisance covariates in each analysis. We note that in such an analysis, there is no inherent need to correct for multiple comparisons since the ability of all individual nodal centrality/ALFF scores to predict group status were tested simultaneously. However, given the large number of potential contributing parameters in the fitted models we chose to retain a subset of results that were the most potent in distinguishing our groups and thus elected p < .005 as a more stringent test of significance.

***Effective Connectivity Amongst Target Regions***

**Rationale and selection of regions for EC analysis.** As noted in the main text, nodal FC analyses revealed hyperconnectivity between daIns and DAN as well as a hypoconnectivity of TPJ in the BPD group. We also found a broad pattern of lower ALFF across a large swath of DAN nodes (see Table S3, Figs S6-7) in the BPD group in addition to higher ALFF in the same daIns node. A large body of research on aIns, TPJ, and DAN indicates that these regions are involved in the dynamic control of attention (30). As such, we sought to understand if the findings from our prior analysis could be integrated to uncover a shared pattern of directed connectivity amongst these regions. Our EC analysis was conceptualized to provide evidence for whether the three salient findings from our whole-brain FC analysis were truly distinct sets of findings or the result of a shared/unified signal. Given previous reports of the EC profile of the insula as a primary controller of other brain regions/networks (i.e., is characterized by a high degree of *output* which feeds into other intrinsic networks), we used LV-GIMME to test directed connectivity amongst DAN, daIns, and TPJ, expecting to find that daIns would be characterized by outgoing edges to DAN.

Results from nodal ALFF and a post-hoc analysis of insula edges indicated that a wide array of DAN regions showed significant group differences (daIns-DAN edges higher and DAN_ALFF_ lower in the BPD group). with no single region clearly driving these results (Tables S4, S6, Figs S6-7). This indicates that DAN hyperconnectivity to daIns and lower ALFF in the BPD group can be better understood as a network level property (defined formally as a latent variable).We selected 15 DAN regions to include as indicators in the estimation of a latent variable in LV-GIMME if they were significant at the p > .005 level in the nodal ALFF analysis (showing lower ALFF) *or* the post-hoc daIns edge analysis (see Table S3). Rather than adding each nodal timeseries as a possible signal to predict/explain in a standard GIMME model, we considered the broad pattern of lower ALFF across spatially contiguous nodes in DAN to be potentially reflective of a shared signal that could be decomposed into a single signal that explains the covariance amongst nodal time series in DAN. We first confirmed the pattern of covariance amongst ALFF in these 15 DAN regions by examining bivariate correlations amongst ALFF scores in these 15 regions (Fig S8). We then fit a single factor model to the set of 15 DAN ALFF scores, and extracted factor scores (31) for use in path models. Factor loadings from the single factor model is presented in Table S6.

**Supplemental Results**

**Global graph characteristics.** Mixed effects regression analyses indicated that groups were not significantly different in global strength centrality (t = 1.16, p = .25) or global ALFF (t = 0.35, p = .73). We found evidence that global ALFF was lower in older adults (t = -2.42, p = .02), with no age-related differences found in strength centrality (t = -0.54, p = .59). Group x age interactions were also nonsignificant in predicting ALFF (t = 0.90, p = .37) and strength centrality (t = -0.62, p = .54). ALFF and strength distributions are presented in Fig S4. We found no significant differences in global graph metrics (Table S2).

| **Supplemental Tables and Figures** | | | | | | |
| --- | --- | --- | --- | --- | --- | --- |
| **Table S1**  *Summary of BPD rsFC studies to date* | | | | | | |
| Study | Activity or connectivity | Analytic technique | Sample size | Age | Results | Notes |
| Wolf et al., (2011) (32) | Functional Connectivity | ICA weights | N_BPD_= 17  N_HC_= 17 | M_BPD_ = 28.6  M_HC_ = 27.2 | ↑ DMN connectivity (L frontopolar cortex, L insula), ↓ R FPN connectivity (IPL, MTG) | R FPN strongly resembles canonical DAN |
| Doll et al., (2013) (33) | Functional Connectivity | ICA (intra-/inter- FC) | N_BPD_= 14  N_HC_= 16 | M_BPD_ = 30.4  M_HC_ = 34.0 | ↑ SN inter-network connectivity to DMN regions | Focus on within- and between-network FC aligns with current study |
| Das et al., (2014) (34) | Functional Connectivity | ICA (inter-FC) | N_BPD_= 14  N_BD_= 16  N_HC_= 13 | M_BPD_ = 32.0  M_BD_ = 35.6  M_HC_ = 31.2 | ↓ SN inter-network connectivity to DMN and RFPN compared to BD patients |  |
| Krause-Utz et al., (2014) (35) | Functional Connectivity | Seed-based (amygdala, dACC) | N_BPD_= 20  N_HC_= 17 | M_BPD_ = 29.6  M_HC_ = 27.5 | ↑ amygdala FC to insula, putamen, OFC, ↓ ACC anticorrelation to PCC |  |
| Baczkowski et al., (2016) (36) | Functional Connectivity | Seed-based (amygdala) | N_BPD_= 48  N_CPD_= 21  N_HC_= 39 | M_BPD_= 30.8  M_CPD_= 31.5  M_HC_= 28.7 | ↓ amygdala FC to PCC, ↑ amygdala FC to SPL and amygdala FC to mPFC, dlPFC, vlPFC unchanged after emotion-regulation task | This study focused on changes in rsFC pre- and post-emotion regulation task |
| Salvador et al., (2016) (37) | Functional Connectivity, Activity | Voxelwise GBC, ALFF | N_BPD_= 60  N_HC_= 60 | M_BPD_ = 32.1  M_HC_ = 33.7 | ↑ ACC and ↓MTG GBC, ↑amygdala/hippocampus and putamen and ↓ Precuneus/ PCC ALFF | This study also found evidence of structural abnormalities in medial structures using DWI |
| Xu et al., (2016) (38) | Functional Connectivity | Graph theory (NBS) | N_BPD_= 20  N_HC_= 10 | M_BPD_ = 29  M_HC_ = 27 | ↑ global graph metrics (small-worldness, local clustering, etc), diffuse NBS network detected with ↓ FC | In our opinion the diffuse nature of metrics and tests under consideration in this study make its findings exceptionally difficult to characterize |
| Lei et al., (2017) (39) | Functional Connectivity, Activity | Seed-based  (R PCC, Precuneus), ALFF, ReHo | N_BPD_= 40  N_HC_= 35 | M_BPD_ = 25.2  M_HC_ = 24.8 | ↓ALFF and ReHo in PCC/Precuneus, ↑ PCC/Precuneus FC to “frontotemporal and limbic lobe” |  |
| Balducci et al., (2018) (40) | Functional Connectivity | Seed-based (amygdala, DMN) | N_BPD+CD+_= 20  N_BPD+CD-_= 10  N_BPD-CD+_ = 19  N_BPD-CD-_= 20 | M_BPD+CD+_ = 31  M_BPD+CD-_ = 31  M_BPD-CD+_ = 31  M_BPD-CD-_ = 32 | BPD+CD+ amygdala-mPFC FC similar to BPD-CD- group. ↓ BPD+CD+ amygdala-insula compared to all other groups | CD = cocaine dependence |
| Lei et al., (2018) (41) | Functional Connectivity, Activity | Graph theory (degree),  fALFF | N_BPD_= 43  N_HC_= 39 | M_BPD_ = 25.2  M_HC_ = 24.8 | ↑ degree in bilateral precuneus, L MTG and AG, ↑ fALFF in occipital regions, and L MTG, ↓ fALFF in R Precuneus and PCC | Degree is computed voxelwise, and thus nearly identical to GBC analyses reported in Salvador et. al., (2016).    Nearly identical demographics between this study and Lei et. al., 2017, leads one to believe this is the same sample, though this is neither confirmed nor denied in the manuscript |
| Wagner et al., (2018) (42) | Functional Connectivity | Seed-based (LC, NCS, SNc) | N_BPD_= 33  N_HC_= 33 | M_BPD_ = 26.7  M_HC_ = 26.4 | ↑LC-ACC and NCS-FPC FC, lack of negative SNc-dlPFC FC |  |
| Duque-Alarcón et al., (2019) (43) | Functional Connectivity | Seed-based (mPFC, ACC, aIns, pINS, precuneus, amygdala, MTG) | N_BPD_= 18  N_HC_= 15 | M_BPD_ = 31.2  M_HC_ = 32.8 | ↑ mPFC-SPL FC, ↓mPFC-Nacc/OFC, ACC-SFG, amygdala-superior parietal, and MTG-motor FC |  |
| Lei et al., (2019) (44) | Functional Connectivity | Seed-based (ACC) | N_BPD_= 43  N_HC_= 39 | M_BPD_ = 25.3  M_HC_ = 24.8 | ↑ L ACC-R MFG, L MTG FC | Similar open questions about the sample used in Lei et. al., 2017, 2018. This study also included DWI analysis |
| Metz et al., (2019) (45) | Functional Connectivity | Seed-based (amygdala, hippocampus) | N_BPD_= 20  N_PTSD_= 18  N_HC_= 40 | M_BPD_ = 27.9  M_PTSD_ = 29.7  M_HC_ = 28.7 | ↓ hippocampus-dmPFC FC, null results for amygdala FC | Included hydrocortisone administration, which did not influence FC of seeds |
| Quattrini et al., (2019) (46) | Functional Connectivity | ICA (intra-FC) | N_BPD_= 21  N_HC_= 14 | M_BPD_ = 35  M_HC_ = 38 | ↓ mean DMN (R PCC focally), SN, and FPN FC |  |
| Reich et al., (2019) (47) | Functional Connectivity | Seed-based (amygdala) | N_BPD_= 14  N_BDII_= 15 | M_BPD_ = 28.1  M_BDII_ = 26.4 | ↓ R amygdala – bilateral MFG FC compared to BDII |  |
| Sarkheil et al., (2019) (48) | Functional Connectivity | ICC (voxelwise avg r^2^),  Seed-based (extracted from ICC) | N_BPD_= 26  N_HC_= 26 | M_BPD_ = 25.2  M_HC_ = 24.9 | ↑ ICC in L aIns, caudate, visual association cortex. ↑ caudate- ACC/mpfc/VS FC and L aIns- midcingulate/R aIns/IPL | VS designation appears to more closely resemble portions of dlS rather than the most ventral portion of the striatum (incl Nacc). |
| Hall & Hallquist, (2022) (49) | Effective Connectivity | Graph theory (GIMME, fronto-limbic nodes) | N_BPD_= 40  N_HC_= 42 | M_BPD_ = 20.8  M_HC_ = 20.6 | ↑ Basolateral amygdala directed influence on central amygdala, ↓ L vmPFC directed influence on L central amygdala | Same sample, parcellation and preprocessing procedure of current study |
| Current study | Functional Connectivity, Effective Connectivity, Activity | Graph theory (network-centrality, LV-GIMME), ALFF | N_BPD_= 40  N_HC_= 42 | M_BPD_ = 20.8  M_HC_ = 20.6 | ↑ R daIns FC to DAN, indicated by ↑ FC between parietal nodes and R daINS, ↓ ALFF across DAN and ↑ ALFF in R daIns, ↑ EC from daIns to TPJ and DAN, EC from daIns to DAN and TPJ |  |
| *Note.* Non-exhuastive search of rs-fMRI studies in borderline patients, conducted on reports published prior to February 2022. | | | | | | |

**Table S2**

*Global Graph Metrics*

| Metric | BPD | HC | Group (p) | Group x Age |
| --- | --- | --- | --- | --- |
| Modularity | 0.038(0.018) | 0.035 (0.016) | 0.30 (0.77) | -0.33 (0.74) |
| Transitivity | 0.928(0.033) | 0.938(0.031) | -1.57 (0.12) | 1.42 (0.16) |
| Global efficiency | 0.240(0.041) | 0.248(0.040) | -1.33 (0.19) | -0.98 (0.33) |
| Characteristic path length | 1.09(0.04) | 1.10(0.04) | 1.23 (0.22) | -1.12 (0.27) |
| Diameter | 16.77(3.12) | 15.71 (3.24) | -0.78 (0.43) | 0.43 (0.67) |
| *Note.* Group and group x age effects for regression models predicting global graph metrics. The first two columns give the mean and SD of the graph metric per group, while the second two columns denote the respective t-scores and p-values for main effects of group and the interaction of group and age in predicting the global metric of interest. All models included the subject’s mean FC as a covariate of no interest. Developmental trajectories are displayed in figure S5. | | | | |

| **Table S3**  *Significant Whole-brain nodal (FC and ALFF) ridge regression effects* | | | | | |
| --- | --- | --- | --- | --- | --- |
| Region/Node (ROI number) | Region Network | Metric | Effect | Est.(S.E.) | t_Ridge_ |
| **R daINS (Roi307)** | **SN** | **NSSC_Vis_** | **Group** | **0.357(0.096)** | **3.70**** |
|  |  | **NSSC_SomMot_** | **Group** | **0.149(0.038)** | **3.93***** |
|  |  | **NSSC_DAN_** | **Group** | **0.016(0.003)** | **4.55***** |
|  |  | **NSSC_SN_** | **Group** | **0.784(0.234)** | **3.35**** |
|  |  | **NSSC_FPN_** | **Group** | **0.241(0.078)** | **3.11*** |
|  |  | **NSSC_DMN_** | **Group** | **0.165(0.050)** | **3.58**** |
|  |  | **ALFF** | **Group** | **0.023(0.006)** | **3.763**** |
| **R TPJ (Roi295)** | **SN** | **Strength** | **Group** | **-0.003(0.001)** | **-2.99*** |
|  |  | **NSSC_Vis_** | **Group** | **-0.288(0.093)** | **-3.11*** |
|  |  | **NSSC_DAN_** | **Group** | **-0.011(0.003)** | **-3.30**** |
|  |  | **NSSC_FPN_** | **Group** | **-0.272(0.073)** | **-3.71**** |
|  |  | **NSSC_DMN_** | **Group** | **-0.143(0.043)** | **-3.31**** |
|  |  | **ALFF** | **Group x Age^2^** | **-0.021(0.006)** | **-3.35**** |
| R Putamen (Roi419) | SN | NSSC_DAN_ | Group x Age | 0.010(0.003) | 2.87* |
| L daINS (Roi102) | SN | NSSC_DMN_ | Group | 0.121(0.042) | 2.89* |
| L anterior temporal gyrus (Roi97) | SN | ALFF | Group | 0.018(0.006) | 2.86* |
| L daINS (Roi103) | SN | ALFF | Group | 0.020(0.006) | 3.13* |
| L dACC (Roi107) | SN | ALFF | Group | 0.022(0.006) | 3.62** |
| R pMTS (Roi294) | SN | ALFF | Group x Age^2^ | -0.019(0.006) | -3.03* |
| R SMG (Roi296) | SN | ALFF | Group x Age^2^ | -0.019(0.006) | -3.01* |
| L SPL (Roi82) | DAN | NSSC_SN_ | Group x Age^2^ | 0.560(0.179) | 3.12* |
| **R Precuneus (Roi288)** | **DAN** | **ALFF** | **Group** | **-0.022(0.007)** | **-3.37**** |
|  |  | **ALFF** | **Group x Age** | **-0.021(0.007)** | **-3.23*** |
| **R Parieto-occip sulcus (Roi277)** | **DAN** | **ALFF** | **Group** | **-0.022(0.006)** | **-3.43**** |
| **R SPL (Roi280)** | **DAN** | **ALFF** | **Group** | **-0.019(0.006)** | **-3.03*** |
| **R SPL (Roi282)** | **DAN** | **ALFF** | **Group** | **-0.020(0.007)** | **-3.09*** |
| **L IPL (Roi73)** | **DAN** | **ALFF** | **Group** | **-0.020(0.006)** | **-3.31**** |
| **L IPL (Roi76)** | **DAN** | **ALFF** | **Group** | **-0.019(0.006)** | **-3.12*** |
| **L SPL (Roi81)** | **DAN** | **ALFF** | **Group** | **-0.022(0.007)** | **-3.37**** |
| R VS (Roi420) | Limbic | NSSC_DMN_ | Group x Age | -0.108(0.038) | -2.84* |
| L VS (Roi415) | Limbic | NSSC_FPN_ | Group x Age^2^ | -0.168(0.057) | -2.96* |
| L Temporal Pole (Roi125) | Limbic | NSSC_Limbic_ | Group x Age | -0.141(0.038) | -3.75** |
| R Inf temporal gyrus (Roi328) | Limbic | ALFF | Group | 0.022(0.007) | -3.26* |
| L mOFC (Roi116) | Limbic | ALFF | Group x Age | -0.019(0.007) | -2.81* |
| R IPL (Roi336) | FPN | NSSC_DAN_ | Group | 0.010(0.003) | 2.90* |
| L Precuneus (Roi144) | FPN | NSSC_Limbic_ | Group x Age | -0.107(0.036) | -2.93* |
| R IPL (Roi333) | FPN | NSSC_FPN_ | Group x Age | 0.201(0.061) | 3.44** |
| R Mid Frontal Gyrus (Roi352) | FPN | NSSC_DMN_ | Group x Age | 0.112(0.037) | 3.00* |
| L IPL (Roi127) | FPN | NSSC_DMN_ | Group x Age | -0.106(0.038) | -2.81* |
| R Sup occipital gyrus (Roi334) | FPN | ALFF | Group x Age | -0.022(0.006) | -3.52* |
| L IFG (Roi136) | FPN | ALFF | Group x Age^2^ | -0.017(0.006) | -2.96* |
| L Sup Frontal Gyrus (Roi186) | DMN | NSSC_SN_ | Group | 0.707(0.231) | 3.06* |
|  |  | NSSC_FPN_ | Group | 0.218(0.077) | 2.84* |
| L Precuneus (Roi200) | DMN | NSSC_Limbic_ | Group | -0.113(0.037) | -3.03* |
|  |  | NSSC_FPN_ | Group x Age | -0.191(0.061) | -3.14* |
| L rACC (Roi174) | DMN | NSSC_SomMot_ | Group | -0.121(0.038) | -3.20* |
| L MTG (Roi157) | DMN | NSSC_Vis_ | Group | -0.288(0.100) | -2.88* |
| R Mid OFC (Roi376) | DMN | NSSC_Vis_ | Group | 0.279(0.099) | 2.83* |
| R Thalamus (Roi412) | DMN | NSSC_FPN_ | Group x Age^2^ | -0.165(0.058) | -2.83* |
| R PCC (Roi398) | DMN | NSSC_DMN_ | Group x Age | -0.118(0.041) | -2.89* |
| L sgACC (Roi169) | DMN | ALFF | Group | 0.022(0.007) | 3.39** |
| R Postcentral Gyrus (Roi253) | SomMot | NSSC_Vis_ | Group | -0.286(0.101) | -2.84* |
| R SMA (Roi254) | SomMot | NSSC_Limbic_ | Group x Age | -0.124(0.038) | -3.28* |
| L Postcentral Gyrus (Roi43) | SomMot | NSSC_Limbic_ | Group x Age^2^ | -0.098(0.034) | -2.89* |
| L Thalamus (Roi406) | SomMot | ALFF | Group | 0.019(0.006) | 3.27* |
| L Mid Occipital Gyrus (Roi14) | Vis | NSSC_Vis_ | Group | 0.348(0.100) | 3.48** |
|  |  | NSSC_SomMot_ | Group | 0.142(0.039) | 3.68** |
|  |  | NSSC_DAN_ | Group | 0.012(0.003) | 3.54** |
|  |  | NSSC_SN_ | Group | 0.670(0.232) | 2.89* |
|  |  | NSSC_FPN_ | Group | 0.258(0.078) | 3.31** |
|  |  | NSSC_DMN_ | Group | 0.177(0.043) | 4.04*** |
| R Cuneus (Roi229) | Vis | NSSC_SomMot_ | Group | 0.117(0.037) | 3.14* |
| R Lingual Gyrus (Roi207) | Vis | NSSC_DAN_ | Group | -0.100(0.003) | -2.84* |
| R Fusiform Gyrus (Roi201) | Vis | NSSC_DAN_ | Group | -0.011(0.003) | -3.06* |
| R Lingual Gyrus (Roi206) | Vis | NSSC_Limbic_ | Group x Age^2^ | -0.094(0.032) | -2.93* |
| R ITG (Roi209) | Vis | NSSC_Limbic_ | Group | -0.122(0.039) | -3.09* |
| L Inf Occipital Gyrus (Roi12) | Vis | NSSC_DMN_ | Group | 0.145(0.044) | 3.29* |
| L Fusiform gyrus (Roi2) | Vis | ALFF | Group x Age | -0.019(0.006) | -3.09* |
| L IPL (Roi31) | Vis | ALFF | Group | -0.018(0.006) | -2.97* |
| *Note.* Nodal ridge regression results that fell below a conservative alpha of .005 (***p < .0001, **p <.001, *p<.005). To aide in visual inspection, we grouped our results into sections. From top to bottom, sections separated by thick bars represent nodes within a given network. Node labels in the first column denote labels taken from Schaefer et. al., (2018) for comparability, while node numbers above 400 were subcortical nodes added from various parcellations (see Method). All metrics marked NSSC_X_ refer to one of seven NSSC scores, denoting normalized strength centrality (overall weighted connectivity) of that individual node with network X (see graph metrics for more details). | | | | | |

| **Table S4**  *Post-hoc analysis of daINS edges* | | | | |
| --- | --- | --- | --- | --- |
| Node (ROI number) | Network | Effect | Est.(S.E.) | t-score |
| R Precuneus (Roi288) | DAN | Group | 0.030(0.007) | 4.37*** |
| L SPL (Roi82) | DAN | Group | 0.026(0.006) | 4.24*** |
| R SPL (Roi282) | DAN | Group | 0.025(0.006) | 3.83** |
| L SPL (Roi81) | DAN | Group | 0.024(0.006) | 3.80** |
| R SPL (Roi289) | DAN | Group | 0.024(0.006) | 3.72** |
| R SPL (Roi286) | DAN | Group | 0.021(0.006) | 3.53** |
| R IPL (Roi283) | DAN | Group | 0.021(0.006) | 3.51** |
| R Postcentral Gyrus (Roi276) | DAN | Group | 0.019(0.006) | 3.17* |
| R Precuneus (Roi285) | DAN | Group | 0.020(0.007) | 3.12* |
| L SPL (Roi84) | DAN | Group | 0.021(0.007) | 3.11* |
| R FEF (Roi291) | DAN | Group | 0.018(0.006) | 2.83* |
| R BLA (Roi 402) | Limbic | Group x Age | -0.020(0.006) | -3.27* |
| L Lingual Gyrus (Roi9) | Vis | Group | 0.023(0.007) | 3.47** |
| L Mid Occipital Gyrus (Roi14) | Vis | Group | 0.020(0.006) | 3.21* |
| R Calcarine Gyrus (Roi215) | Vis | Group | 0.019(0.006) | 3.02* |
| R Inf Occipital Gyrus (Roi213) | Vis | Group | 0.020(0.007) | 3.00* |
| R Sup Occipital Gyrus (Roi230) | Vis | Group | 0.017(0.006) | 2.90* |
| L Inf Occipital Gyrus (Roi12) | Vis | Group | 0.019(0.007) | 2.82* |
| R Sup Occipital Gyrus (Roi221) | Vis | Group | 0.018(0.006) | 2.81* |
| R Calcarine Gyrus (Roi224) | SomMot | Group | 0.021(0.006) | 3.28* |
| L Precentral Gyrus (Roi65) | SomMot | Group | 0.020(0.006) | 3.18* |
| L Heschl’s Gyrus (Roi34) | SomMot | Group x Age | -0.019(0.006) | -3.13* |
| R MCC (Roi250) | SomMot | Group | 0.019(0.006) | 3.04* |
| R STG (Roi231) | SomMot | Group | 0.019(0.007) | 2.90* |
| R Postcentral Gyrus (Roi267) | SomMot | Group | 0.019(0.007) | 2.86* |
| R Angular Gyrus (Roi334) | FPN | Group | 0.018(0.006) | 2.89* |
| *Note*. Significant results (***p < .0001, **p <.001, *p<.005) from a post-hoc analysis of all edges incident to daIns. Higher estimates denote edge values that are higher in the BPD group. | | | | |

**Table S5**

*Selected L2 penalty (λ) for ridge regressions*

| Analysis | λ |
| --- | --- |
| Strength | 1.99 |
| Vis_z | 0.001 |
| SomMot_z | 6.51 |
| DAN_z | 70.02 |
| Sal_z | 0.22 |
| Limbic_z | 8.28 |
| FPN_z | 3.64 |
| DMN_z | 7.93 |
| ALFF | 32.25 |
| Post-hoc daIns edges | 0.33 |
| *Note.* Ridge penalty parameter derived using automated algorithm described in (27) and implemented in R *ridge* package (28) | |

**Table S6**

*DAN Nodes Selected for LV-GIMME and Path Models*

| Node (ROI name) | FC to R daIns | FC to R TPJ | ALFF | Factor Loading |
| --- | --- | --- | --- | --- |
| L IPL (Roi73) | 0.21 | -2.44 | **-3.31**** | 0.73 |
| L IPL (Roi76) | 0.74 | 1.10 | **-3.12*** | 0.76 |
| L SPL (Roi81) | **3.80**** | 1.06 | **-3.37**** | 0.70 |
| L SPL (Roi82) | **4.24***** | -1.11 | -1.51 | 0.82 |
| L SPL (Roi84) | **3.11*** | -0.05 | -1.83 | 0.74 |
| R Postcentral Gyrus (Roi276) | **3.17*** | 0.58 | 0.94 | 0.65 |
| R Parieto-occip sulcus (Roi277) | 2.80 | -0.33 | **-3.43**** | 0.73 |
| R SPL (Roi280) | 1.29 | -1.41 | **-3.03*** | 0.78 |
| R SPL (Roi282) | **3.83**** | -0.07 | **-3.09*** | 0.77 |
| R IPL (Roi283) | **3.51**** | 0.79 | -0.46 | 0.80 |
| R Precuneus (Roi285) | **3.12*** | -0.14 | -1.80 | 0.69 |
| R SPL (Roi286) | **3.53**** | **-3.05*** | -2.03 | 0.80 |
| R Precuneus (Roi288) | **4.37***** | 0.46 | **-3.37**** | 0.72 |
| R SPL (Roi289) | **3.72**** | -1.89 | -0.82 | 0.71 |
| R FEF (Roi291) | **2.83*** | 0.46 | 0.35 | 0.53 |
| *Note.* Test statistics (t) and significance levels (***p < .0001, **p <.001, *p<.005) for three logistic ridge regression models fit to a) all undirected edges incident to R daIns, b) all edges incident to R TPJ, and c) all nodal ALFF scores. Test statistics are extracted for those parameters in the logistic model that correspond to one of the 15 DAN nodes that were selected for dimension reduction in further targeted analyses. Higher test statistics indicate that the value is higher in the BPD group, with negative statisitics denoting lower values in the BPD group. Factor loadings are taken from a single factor EFA used to extract factor scores for inclusion in path models as an aggregate measure of DAN ALFF. | | | | |

| **Table S7**  *Linear regression on LV-GIMME edge values* | | | | |
| --- | --- | --- | --- | --- |
| Outcome | Predictor | Est. (S.E.) | *t* | *p* |
| daIns→DAN_EC_ | **Group** | **0.07(0.01)** | **5.27** | **< .001** |
|  | Age | 0.02(0.01) | 1.84 | .07 |
|  | $\bar{\mathbf{FC}}$ | **0.40(0.15)** | **2.63** | **.01** |
|  | Group x Age | -0.02(0.01) | 1.36 | .18 |
| daIns→TPJ_EC_ | **Group** | **0.04(0.02)** | **2.04** | **.04** |
|  | Age | 0.02(0.01) | 1.092 | .28 |
|  | $\bar{\mathbf{FC}}$ | **1.22(0.23)** | **5.43** | **< .001** |
|  | Group x Age | 0.00(0.02) | -0.03 | .97 |
| TPJ→DAN_EC_ | **Group** | **-0.11(0.02)** | **-6.15** | **< .001** |
|  | Age | -0.00(0.01) | -0.32 | .75 |
|  | $\bar{\mathbf{FC}}$ | **0.66(0.20)** | **3.31** | **< .01** |
|  | Group x Age | 0.01(0.02) | -0.35 | .73 |
| Note. Results from three linear regressions separated by thin lines. Higher values for group predictor denotes EC edges higher in BPD group. Average whole-brain FC value per subject was included as a nuisance covariate. | | | | |

| **Table S8**  *Parameter table: daIns→DAN Model* | | | | | | |
| --- | --- | --- | --- | --- | --- | --- |
| Parameter Type | Outcome | Predictor | *B* | Lower CI (2.5%) | Upper CI (97.5%) | *p* (two-tailed) |
| Regression | DAN_ALFF_ | **daIns_ALFF_** | **-0.110** | **-0.221** | **0.000** | **0.050^*^** |
|  |  | **daIns→DAN_EC_** | **-0.891** | **-1.816** | **0.050** | **0.064^†^** |
|  |  | daIns**↔︎**DAN_FC_ | -0.10 | -0.098 | 0.078 | 0.832 |
|  |  | $\bar{\text{FC}}$ | 0.960 | -0.724 | 2.623 | 0.266 |
|  |  | $\bar{\text{ALFF}}$ | **0.360** | **0.270** | **0.450** | **<0.001^***^** |
|  | daIns→DAN_EC_ | **daIns_ALFF_** | **0.046** | **0.018** | **0.072** | **<0.001^***^** |
|  |  | $\bar{\text{FC}}$ | **0.473** | **0.053** | **0.902** | **0.028^*^** |
|  |  | $\bar{\text{ALFF}}$ | **-0.032** | **-0.054** | **-0.009** | **0.006^**^** |
|  | daIns**↔︎**DAN_FC_ | daIns_ALFF_ | 0.237 | -0.060 | 0.534 | 0.120 |
|  |  | $\bar{\text{FC}}$ | -3.194 | -7.768 | 1.387 | 0.164 |
|  |  | $\bar{\text{ALFF}}$ | 0.063 | -0.179 | 0.304 | 0.606 |
| Connectivity Correlation | **daIns→DAN_EC_** | **daIns↔︎DAN_FC_** | **0.020** | **0.009** | **0.036** | **<0.001^***^** |
| Residual Variances | **DAN_ALFF_** |  | **0.067** | **0.049** | **0.094** | **<0.001^***^** |
|  | **daIns→DAN_EC_** |  | **0.005** | **0.004** | **0.007** | **<0.001^***^** |
|  | **daIns↔︎DAN_FC_** |  | **0.560** | **0.414** | **0.792** | **<0.001^***^** |
| Intercepts | **DAN_ALFF_** |  | **0.775** | **0.308** | **1.244** | **0.002^***^** |
|  | **daIns→DAN_EC_** |  | **-0.127** | **-0.249** | **-0.007** | **0.040^*^** |
|  | daIns**↔︎**DAN_FC_ |  | -0.833 | -2.156 | 0.492 | 0.210 |
| Mediation: Indirect Effects | **daIns_ALFF_ → daIns→DAN_EC_ → DAN_ALFF_** | | **-0.038** | **-0.098** | **0.002** | **0.066^†^** |
|  | daIns_ALFF_ **→** daIns**↔︎**DAN_FC_ **→** DAN_ALFF_ | | -0.001 | -0.031 | 0.023 | 0.852 |
| *Note.* CFI = 0.993, TLI = 0.957, RMSEA = 0.10, Posterior Predictive P-Value = 0.462. Est denotes unstandardized Bayesian parameter estimates. Standardized estimates for significant paths are graphically displayed in Fig 2a. | | | | | | |

| **Table S9**  *Parameter table: TPJ→DAN Model* | | | | | | |
| --- | --- | --- | --- | --- | --- | --- |
| Parameter Type | Outcome | Predictor | Est. | Lower CI (2.5%) | Upper CI (97.5%) | *p* (two-tailed) |
| Regression | DAN_ALFF_ | TPJ_ALFF_ | 0.035 | -0.045 | 0.133 | 0.376 |
|  |  | **TPJ→DAN_EC_** | **0.892** | **0.145** | **1.655** | **0.020^*^** |
|  |  | TPJ**↔︎**DAN_FC_ | 0.013 | -0.066 | 0.092 | 0.732 |
|  |  | $\bar{\text{FC}}$ | -0.302 | -2.004 | 1.355 | 0.716 |
|  |  | $\bar{\text{ALFF}}$ | **0.312** | **0.230** | **0.395** | **<0.001^***^** |
|  | TPJ→DAN_EC_ | **TPJ_ALFF_** | **0.043** | **0.018** | **0.067** | **<0.001^***^** |
|  |  | $\bar{\text{FC}}$ | **0.968** | **0.408** | **1.534** | **0.002^***^** |
|  |  | $\bar{\text{ALFF}}$ | **-0.033** | **-0.061** | **-0.005** | **0.022^*^** |
|  | TPJ**↔︎**DAN_FC_ | **TPJ_ALFF_** | **0.555** | **0.314** | **0.796** | **<0.001^***^** |
|  |  | $\bar{\text{FC}}$ | 4.358 | -1.150 | 9.793 | 0.118 |
|  |  | $\bar{\text{ALFF}}$ | -0.191 | -0.467 | 0.083 | 0.170 |
| Connectivity Correlation | TPJ→DAN_EC_ | TPJ**↔︎**DAN_FC_ | **0.043** | **0.024** | **0.069** | **<0.001^***^** |
| Residual Variances | DAN_ALFF_ |  | **0.066** | **0.048** | **0.093** | **<0.001^***^** |
|  | TPJ→DAN_EC_ |  | **0.008** | **0.006** | **0.012** | **<0.001^***^** |
|  | TPJ**↔︎**DAN_FC_ |  | **0.800** | **0.590** | **1.133** | **<0.001^***^** |
| Intercepts | DAN_ALFF_ |  | **0.490** | **0.045** | **0.938** | **0.032^*^** |
|  | TPJ→DAN_EC_ |  | **-0.254** | **-0.398** | **-0.111** | **<0.001^***^** |
|  | TPJ**↔︎**DAN_FC_ |  | **-2.549** | **-3.974** | **-1.129** | **<0.001^***^** |
| Mediation: Indirect Effects | **TPJ_ALFF_ → TPJ→DAN_EC_ → DAN_ALFF_** | | **0.036** | **0.005** | **0.083** | **0.022^*^** |
|  | TPJ_ALFF_ → TPJ**↔︎**DAN_FC_ → DAN_ALFF_ | | 0.007 | -0.038 | 0.053 | 0.730 |
| *Note.* CFI = 0.995, TLI = 0.971, RMSEA = 0.098, Posterior Predictive P-Value = 0.472. Est denotes unstandardized Bayesian parameter estimates. Standardized estimates for significant paths are graphically displayed in Fig 2b. | | | | | | |

**Figure S1**

*Similarity matrix of subject adjacency matrices across subjects*

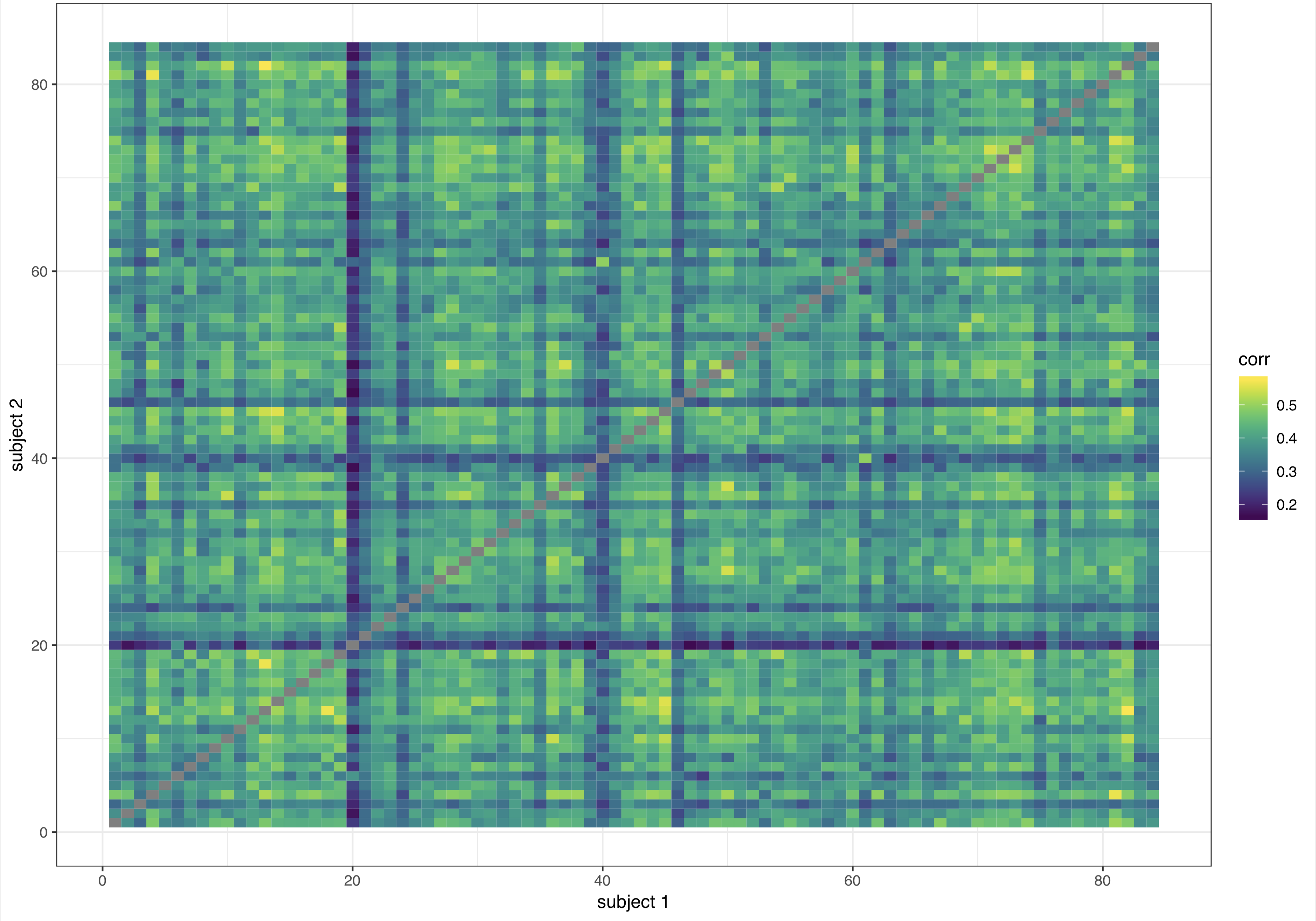

*Note.* Static pearson correlations of individual adjacency matrices by subject. This was conducted on the original 84 subjects that passed our excessive motion screen and is simply meant to demonstrate thinking regarding removal of one subject in the BPD group (who can be spotted by eye in row/column 20) whose adjacency matrix had an unusually low correlation on the whole (~.1) with all other subjects in the study.

**Figure S2**

*Anatomical depiction of custom nodal parcellation*

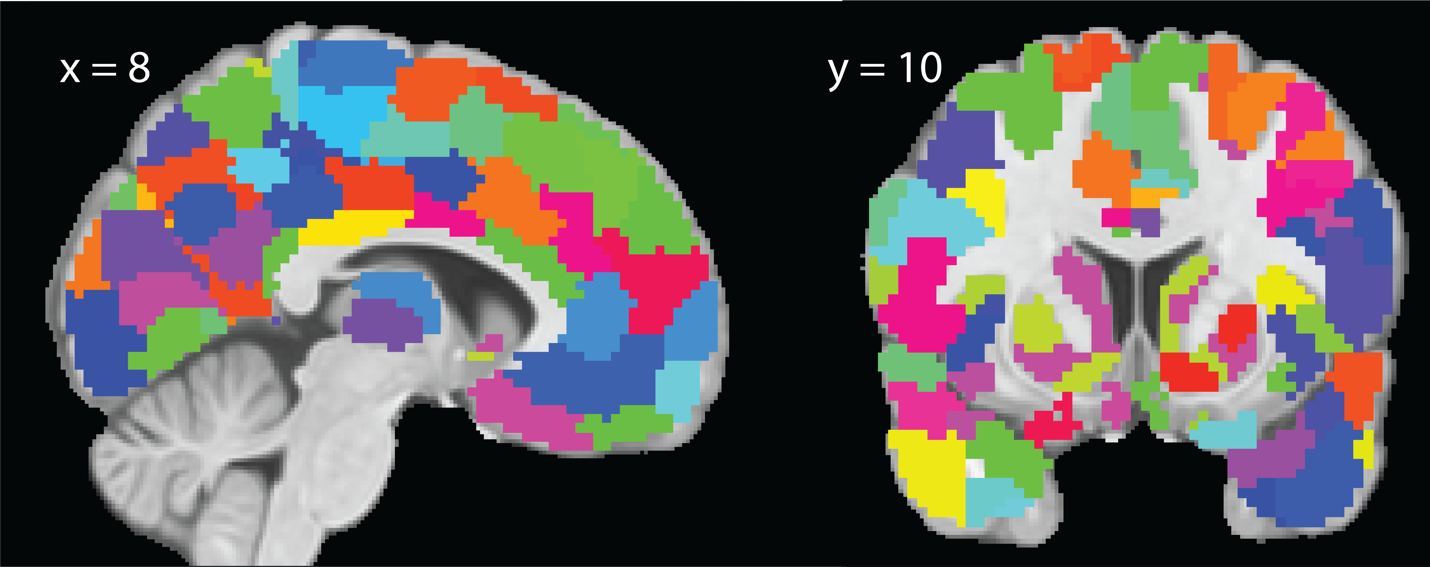

*Note.* Finalized 421-node parcellation of the cortex, thalamus, and striatum from two representative slices. Notice the truncation in inferior orbitofrontal/temporal regions due to significant signal dropout in these voxels.

**Figure S3**

*Average FC within and between intrinsic networks*

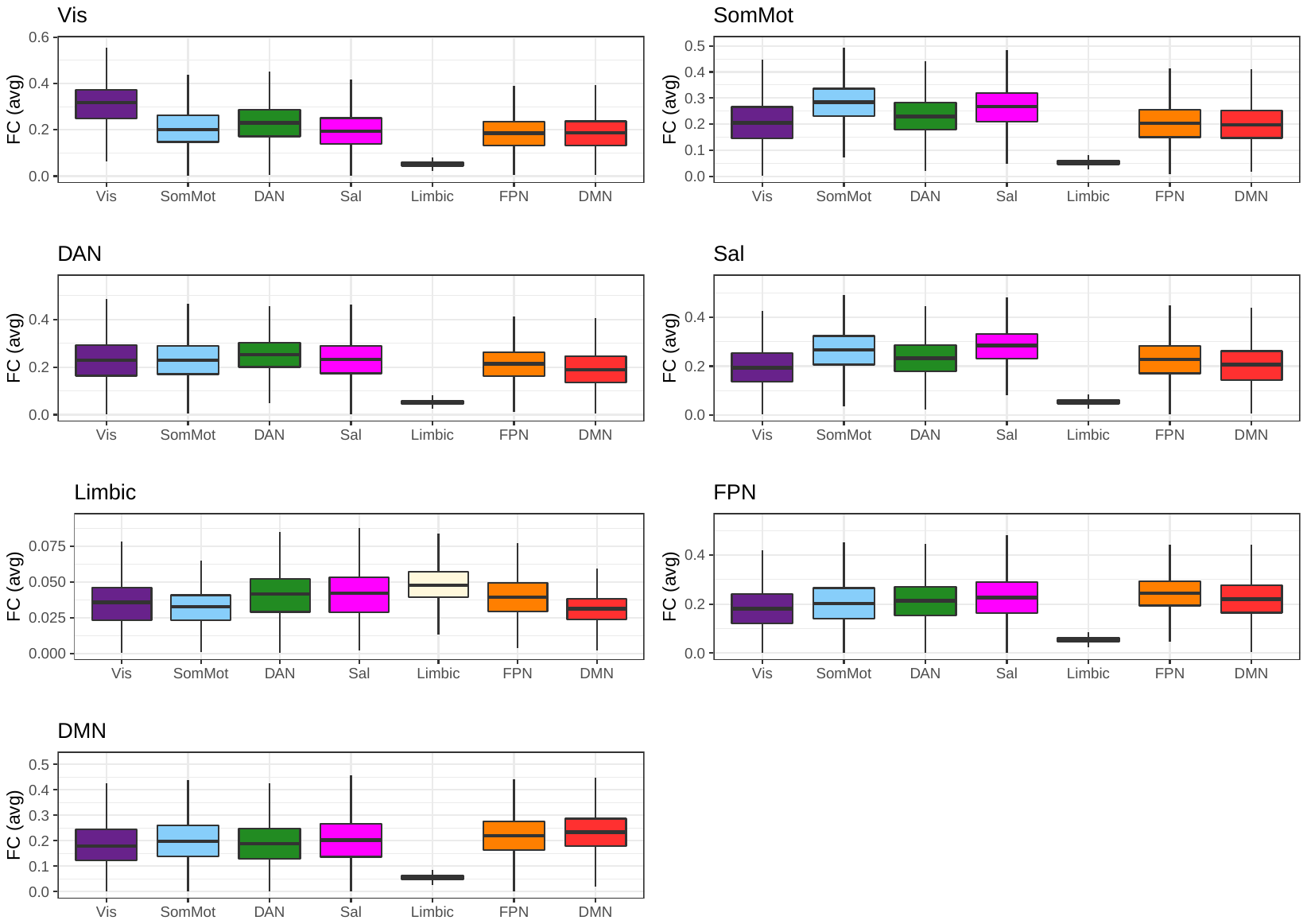

*Note.* In each panel we took all edges from nodes in the corresponding and calculated the average FC value of nodes connecting all seven networks. As expected, edges going from nodes in an intrinsic network to other nodes within the network were generally higher than edges going to nodes that were assigned to other networks.

**Figure S4**

*Strength Centrality and ALFF Distributions Between Groups*

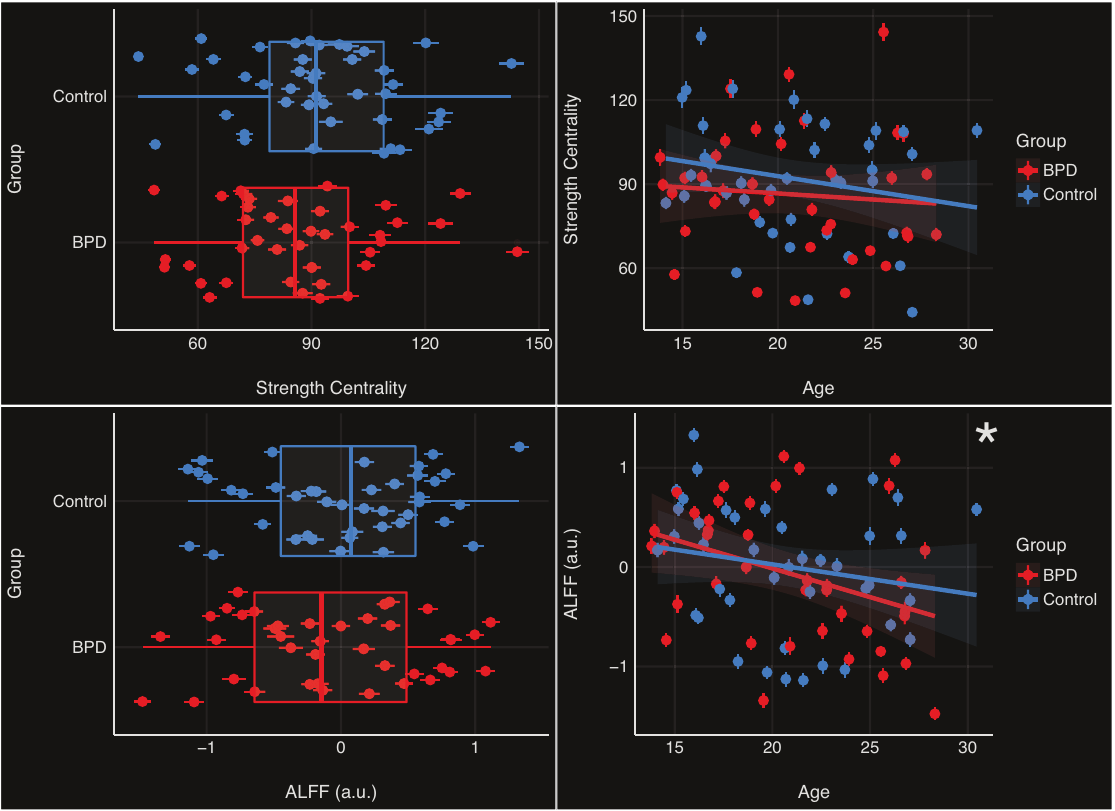

*Note.* Top left panel: Strength centrality (weighted degree) distributions between BPD and Control group. Points represent the median point estimate per subject, whereas error bars expand around the central point estimate ± 2*S.E. Top right panel: Strength centrality as a function of age, error bars around point estimates are identical to those in the left panel. Bottom left panel: ALFF distributions between BPD and Control group. Points represent the median point estimate per subject, whereas error bars expand around the central point estimate ± 2*S.E. Bottom right panel: ALFF as a function of age, error bars around point estimates are identical to those in the left panel. Mixed effects analyses indicated that global ALFF levels linearly decrease with age, with no differences between groups.

**Figure S5**

*Age-related changes in global graph metrics*

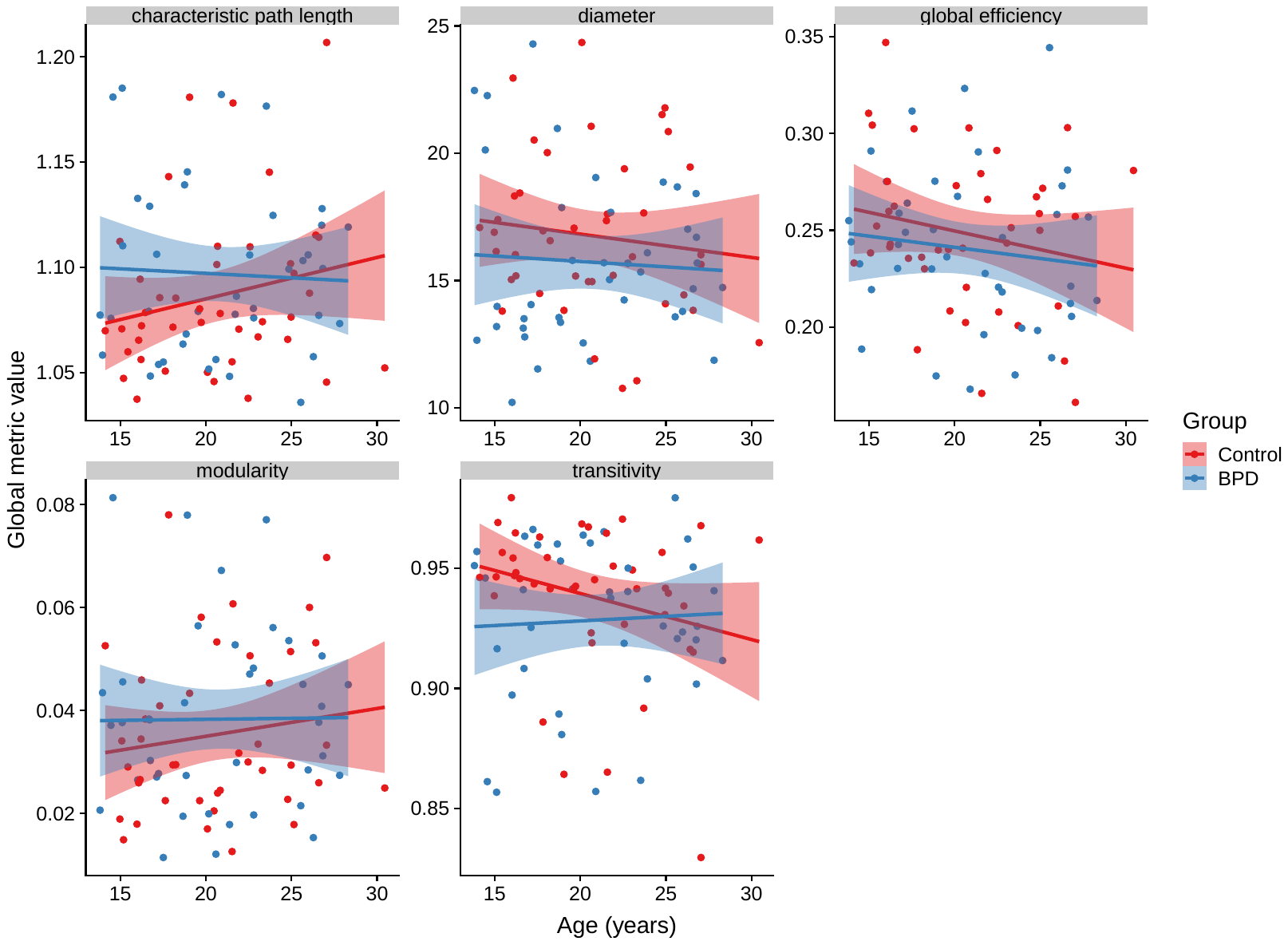

*Note.* All group and group x age effects on global graph metrics were non-significant.

**Figure S6**

*Heatmaps of NSSC (FC) and ALFF analyses: ridge regression effects in model space*

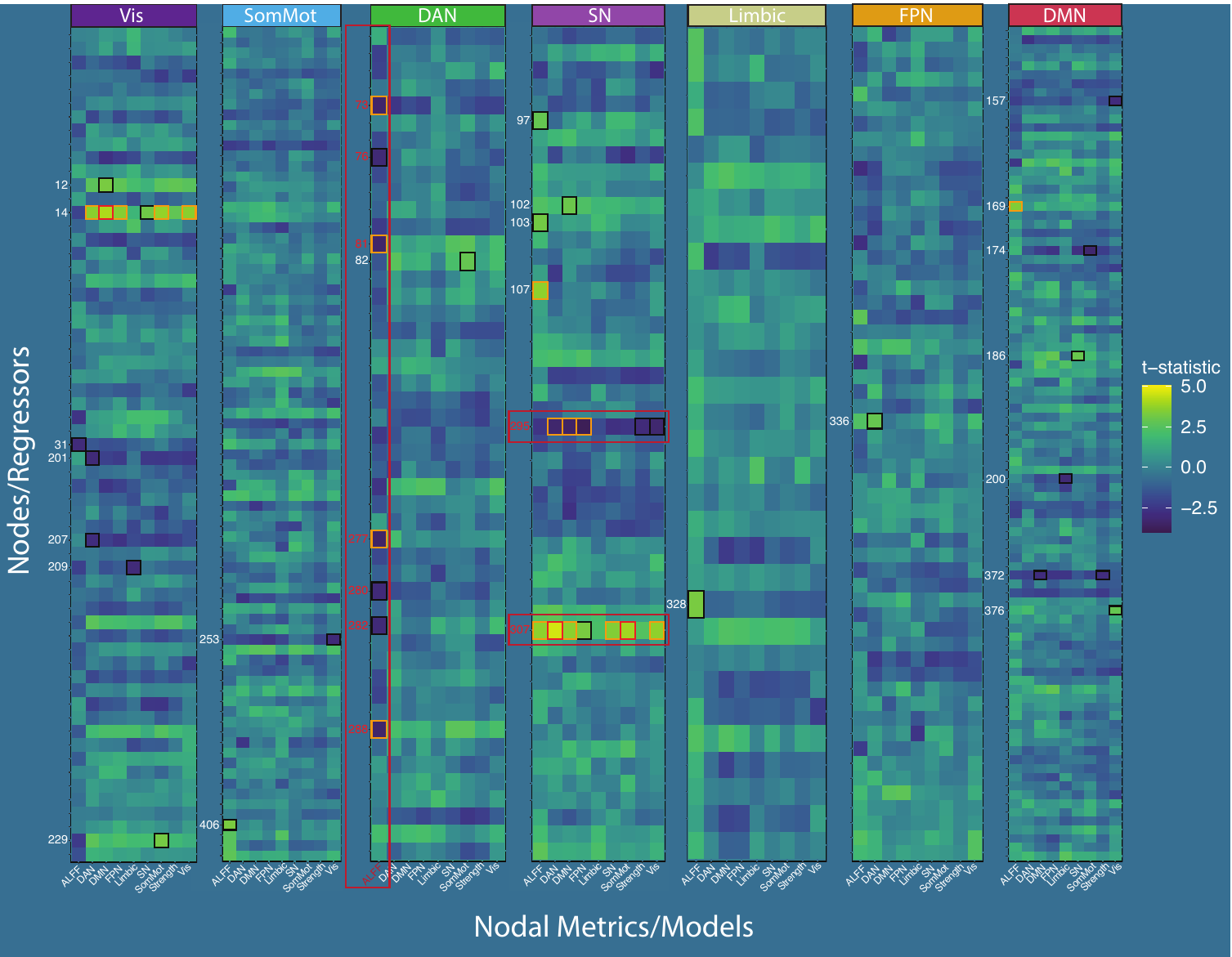

*Note.* An overview of group effects (BPD – HC) within the overall model space. All regions/nodes from our combined parcellation are depicted as rows, which served as joint predictors of group status in a series of logistic ridge regression analyses (run for each nodal metric: strength centrality, seven NSSC scores, and ALFF [depicted as columns]). Cell values denote the parameter t-statistic (higher = heightened in BPD group). At a conservative alpha level of .005, cells displaying significant group differences are highlighted with a black (p < .005), orange (p < .001), or red (p < .0001) borders. For visualization purposes, we grouped nodes/predictors within a given network to a panel of all nodes in that network. Note however, that models were fit jointly across all nodes, meaning for example that all cells in the ALFF (far-left) columns across panels were extracted from a single model rather than being fit separately. Also note that node x age interactions and covariates of no interest (mean FC, mean FD, income level) were included in these models but are not depicted here. We draw the reader’s attention to three key effects: the robust patterns of hyper- and hypo-connectivity across connectivity metrics in the R daIns (307) and R TPJ (295), respectively and the widespread lowered ALFF scores across multiple DAN nodes depicted in the first column of the DAN panel. Full details on all significant effects can be found in Table S2.

**Figure S7**

*DAN nodes show broad pattern of heightened FC to daIns in addition to lower ALFF*

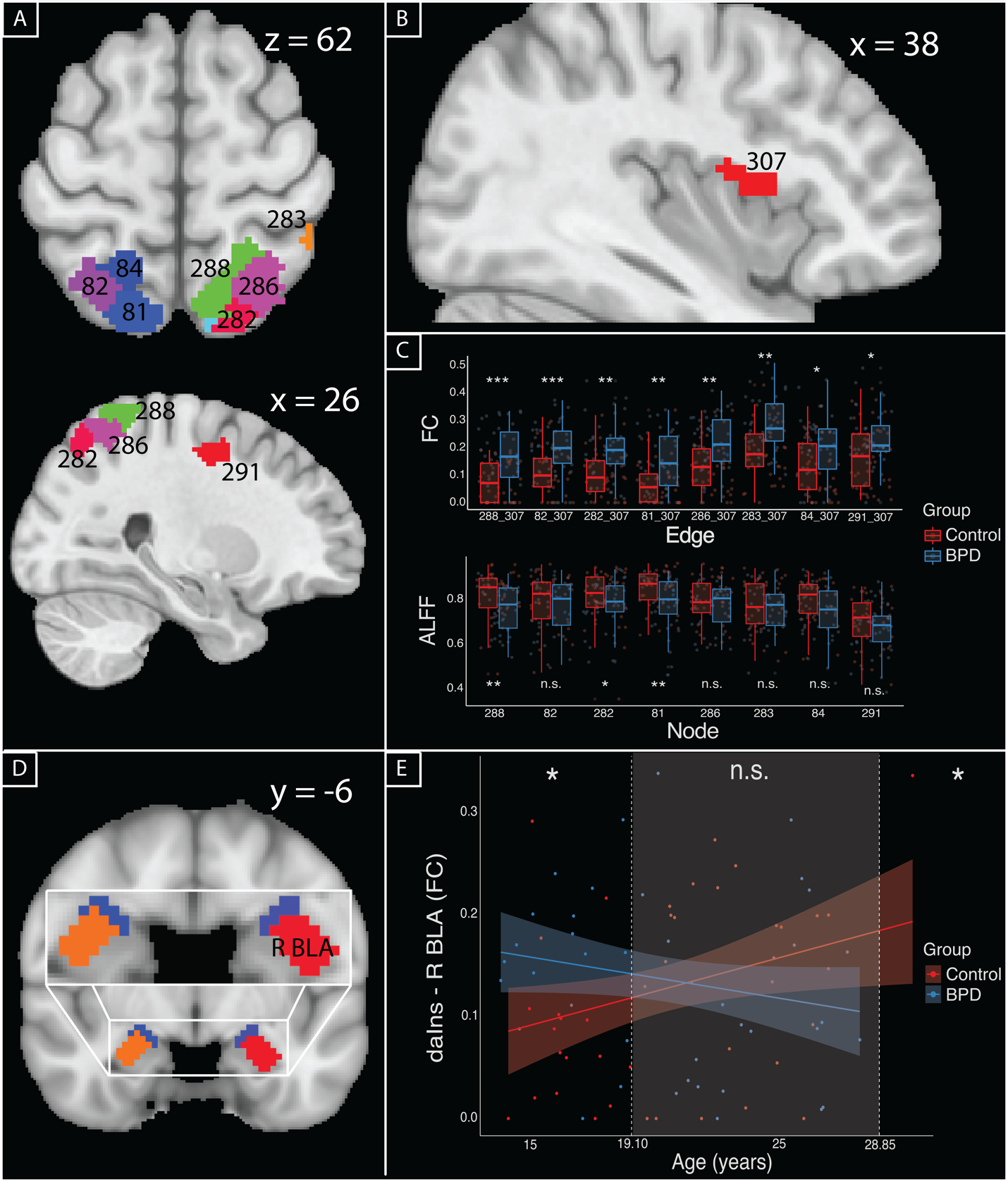

*Note.* (A) Anatomical rendering of 8 representative DAN nodes in MNI space consisting of bilateral SPL and right FEF that showed evidence of hyperconnectivity to the R daIns (B) in edge-focused analysis. (C) Boxplots of the raw edge values and ALFF scores split by group (*p < .005, **p < .001, ***p < .0001).

**Figure S8**

*Bivariate Correlations Amongst ALFF values amongst DAN and SN nodes*
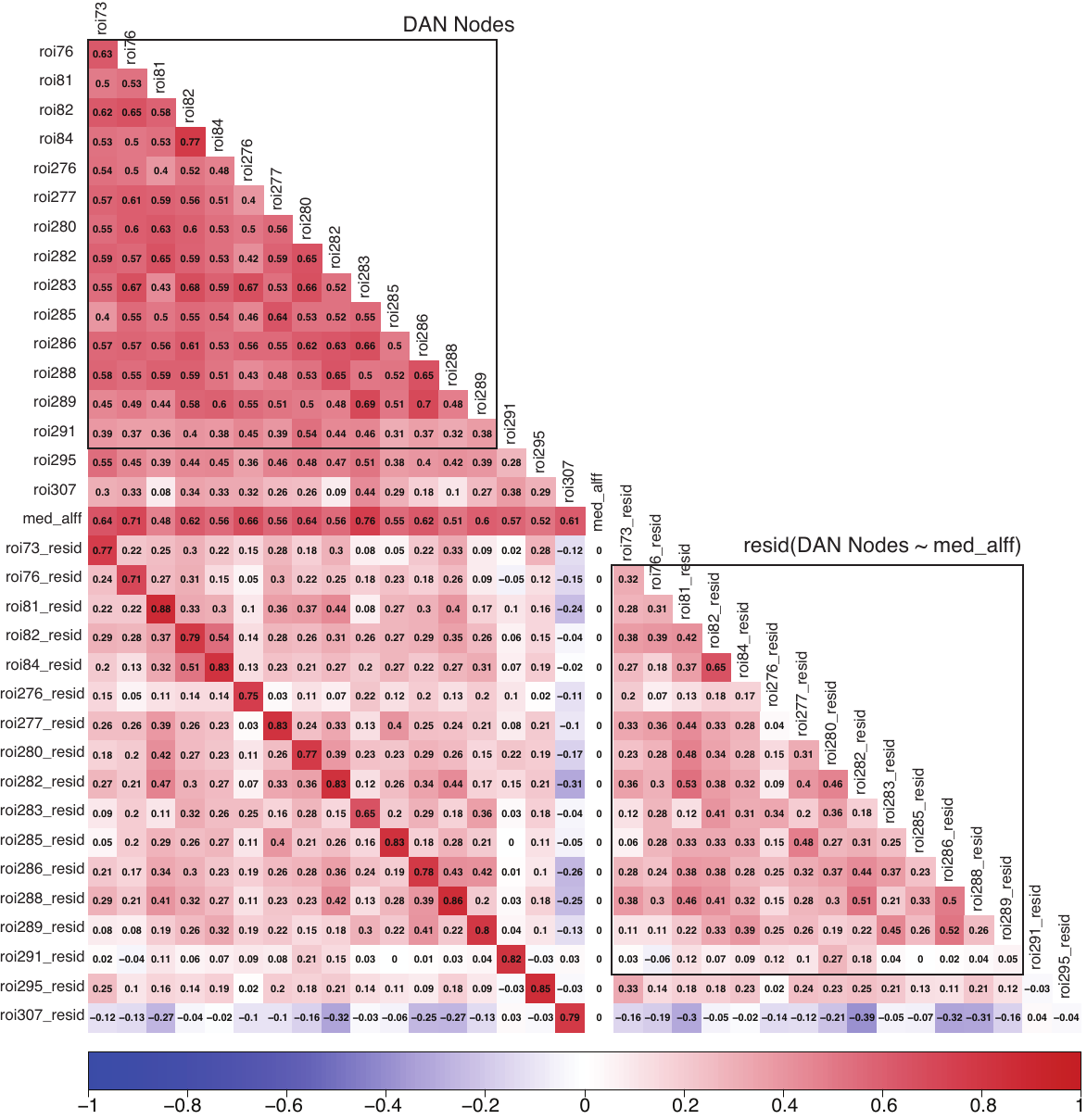

*Note.* Bivariate Pearson correlations amongst ALFF values amongst DAN regions (roi73-291), TPJ(roi295), and daIns (roi307). To show that subject-wise median ALFF (med_alff) induces false positive correlations between DAN nodes and daIns in the low	er right corner, we report correlations of residuals of ALFF values when subject median is regressed out (roiX_resid). Note that correlations between DAN nodes pre-residualizing are positively associated with daIns ALFF, though after residualizing, associations are negative.

1. We elected to compute ALFF on the preprocessed nodal time series prior pre-whitening because ARMA filtering necessarily affects the spectral properties of the nodal time series.
